# Monocytic fibrocyte-like cell enrichment and myofibroblastic adaptation causes nucleus pulposus fibrosis and associates with disc degeneration severity

**DOI:** 10.1101/2024.01.16.575808

**Authors:** Yi Sun, Yan Peng, Zezhuo Su, Kyle KH So, Qiu-ji Lu, Mao-jiang Lyu, Jianwei Zuo, Yong-can Huang, Zhi-ping Guan, Kenneth MC Cheung, Zhao-min Zheng, Xin-tao Zhang, Victor YL Leung

## Abstract

Fibrotic remodeling of nucleus pulposus (NP) leads to structural and mechanical anomalies of intervertebral discs that prone to degeneration, leading to low back pain incidence and disability. Emergence of fibroblastic cells in disc degeneration has been reported, yet their nature and origin remain elusive. In this study, we performed an integrative analysis of multiple single-cell RNA sequencing datasets to interrogate the cellular heterogeneity and fibroblast-like entities in degenerative human NP specimens. We found that disc degeneration severity is associated with an enrichment of fibrocyte-like phenotype, characterized by CD45 and collagen I dual positivity, and expression of myofibroblast marker α-smooth muscle actin. Refined clustering and classification distinguished the fibrocyte-like populations as subtypes in the NP cells - and immunocytes-clusters, expressing disc degeneration markers *HTRA1* and *ANGPTL4* and genes related to response to TGF-β. In injury-induced mouse disc degeneration model, fibrocyte-like cells were found recruited into the NP undergoing fibrosis and adopted a myofibroblast phenotype. Depleting the fibrocyte-like cells in CD11b-DTR mice in which monocytic lineages were ablated by diphtheria toxin could markedly attenuate fibrous modeling and myofibroblast formation in the NP of the degenerative discs, and prevent disc height loss and histomorphological abnormalities. Marker analysis supports that disc degeneration progression is dependent on a function of CD45^+^COL1A1^+^ and αSMA^+^ cells. Our findings reveal that fibrocyte-like cells play a pivotal role in NP fibrosis and may therefore be a target for modifying disc degeneration and promoting its repair.

## Introduction

The intervertebral discs (IVDs) are the largest avascular connective tissues in the body, playing a central role in spine movement. IVD degeneration (IDD) is associated with fibrosis of the nucleus pulposus (NP), the innermost colloidal core of IVDs, along with a loss of hydration and swelling pressure. The degeneration involves a degradation of hyaline extracellular matrix (ECM), in particular aggrecan and collagen II, and accumulation of fibrotic components including collagen I and small leucin-rich proteoglycans such as biglycan and fibronectin^1^. This leads to reduced tissue mechanical strength and hence disc prolapse under load, ultimately causing back pain and myelopathy ^2^. Previous study has shown that transplantation of mesenchymal stromal cells (MSCs)^3^ can alleviate IDD via inhibiting NP fibrosis. Understanding and controlling the profibrotic events may therefore hold the key to modifying the degeneration process. However, the mechanism of NP fibrosis and the origin of the fibroblastic population in the IDD remains elusive.

The primitive NP is initially populated by forkhead transcription factor (*FOXA2*)-or brachyury (*TBXT*)-expressing vacuolated cells derived from the notochord^4–6^. In humans, these vacuolated cells are gradually replaced by the chondrocyte-like cells after birth^7^. Tissue fibrosis relates to an excessive fibroblast and myofibroblast activity ^8^. Reports have shown an increased number of αSMA^+^/FSP1^+^/collagen I^+^ fibroblastic cells in the degenerative IVDs^4,9^. In fact, NP cells (NPC) have a capacity to become FSP1^+^ cells when exposed to excessive TGF-β^5,10^ as well as undergo fibroblastic/myofibroblastic transition (αSMA^+^/COL1A1^+^/FSP1^+^/FAP-α^+^) in an injury-induced IDD model^4^. However, tracing study indicated that not all myofibroblastic cells are derived from local cells^4^, implying alternative origins of effector cells for NP fibrosis. In line with this notion, single cell transcriptomic studies have revealed that the NP of human degenerative IVDs^7,11–14^ and animal model of IDD^6^ contain cell types other than IVD cells. These include endothelial cells, myeloid granulocytic suppressor cells, CCR2^+^ monocytes and derivatives of macrophages. Consistent with the finding, studies showed that bone marrow cells could migrate into mechanically overloaded IVD explants^15^ and that focal enrichment of fibroblastic cells could be found near the infiltrating blood vessels in the degenerative IVDs^16^. These evidences support that access of extrinsic cells to the degenerated IVD may mediate NP fibrosis.

We hypothesized that cell sources other than resident disc cells can contribute to NP fibrosis. In this study, we interrogated the cell hierarchy in the NP of degenerative IVDs through an integrative analysis of the published single cell transcriptomes and characterization in human tissue and cells. Our findings indicated an emergence of a small hematopoietic population that possess a fibrocyte-like phenotype and myofibroblastic identity in the degenerative samples. Using a mouse model that labels or depletes monocyte-derived cells, we analyzed the time course of the disc fibrocytes appearance and identified their role in NP fibrosis and IDD progression. Our study implies a function of circulatory fibrocytes in NP fibrosis and as a target for modifying IDD.

## Results

### Identification of myofibroblastic NPC in disc degeneration and NP fibrosis

We examined the cellular heterogeneity and fibroblastic cell entities by an integrative analysis of 4 published single-cell RNA sequencing (scRNA-seq) data derived from cells isolated from NP tissues^7,12,14^ or recognized as NPC ^11^. The datasets consist of 6 non-degenerative human IVDs, 13 mildly (Pfirrmann II-III) and 12 severely (Pfirrmann IV-V) degenerative human IVDs (**Table 1**). A number of existing NPC phenotypes were previously identified from these datasets, and their marker genes were summarized in **Supplementary Table 1**. The meta-analysis identified 12 cell clusters from a total of 192,184 cells and the cell types were annotated based on their cluster differentially expressed genes (DEGs, Table SI). In addition to 5 NPC clusters, we also identified neutrophils (*LYZ+*), myeloid derived suppressor cells (MDSC+; *MPO+*), macrophages (*CD68+*), T cells (*TRBC2+*), endothelial cells (*CD34+* and *PECAM1+*), pericytes (*RGS5+* and *ACTA2+*), and erythrocytes (*HBB+*) (**Fig. 1a** and **Supplementary Figure S1**). These clusters were mostly present across the four datasets although their proportion and distribution varied (**Fig. 1b**). Clustered NPC were identified by their expression of *ACAN*, *COL2A1* and *SOX9* (Table SI). The most abundant NPC population was the chondrogenic cluster (ChonNP) (51%) which expressed *COL2A1*, *ACAN* and *CYR61*. The regulatory cluster (RegNP) (21%) expressed *MMP3*, *GPX3* and *CHI3L2/1*. These RegNP gene markers were also expressed in the inflammatory C1/3 chonNPC and regulatory NPC reported by Han et al^7^ and Tu et al^12^, respectively (**Supplementary Table S1**). The fibroblastic cluster (FibroNP; 8%) expressed *COL1A1*, *COL3A1*, *POSTN* and *ASPN*, which are known to associate with tissue fibrosis. We found that the proportion of FibroNP significantly increased with the severity of disc degeneration and became one of the major NPC populations (30%) in severely degenerated NP (**Fig. 1c and Supplementary Figure S2**). The cycling cluster (CyclingNP) (1%) expressed the cell proliferation genes including *TOP2A* and *STMN1*, containing two distinct subsets expressing high level of either collagen I or II gene (*COL1A1^high^* and *COL2A1^high^*). The progenitor cluster (ProgNP) (1%), distinguished by the expression of *PDGFRA* and *PLA2G2A*, was unique to the non-degenerative samples as reported by Gan et al^14^. Endothelial cells (2%) were also found, which highly expressed *PECAM1* and *CD34*. Erythrocytes (1%) were defined by gene expression of *HBB* (**Fig. 1d**).

**Figure 1.**
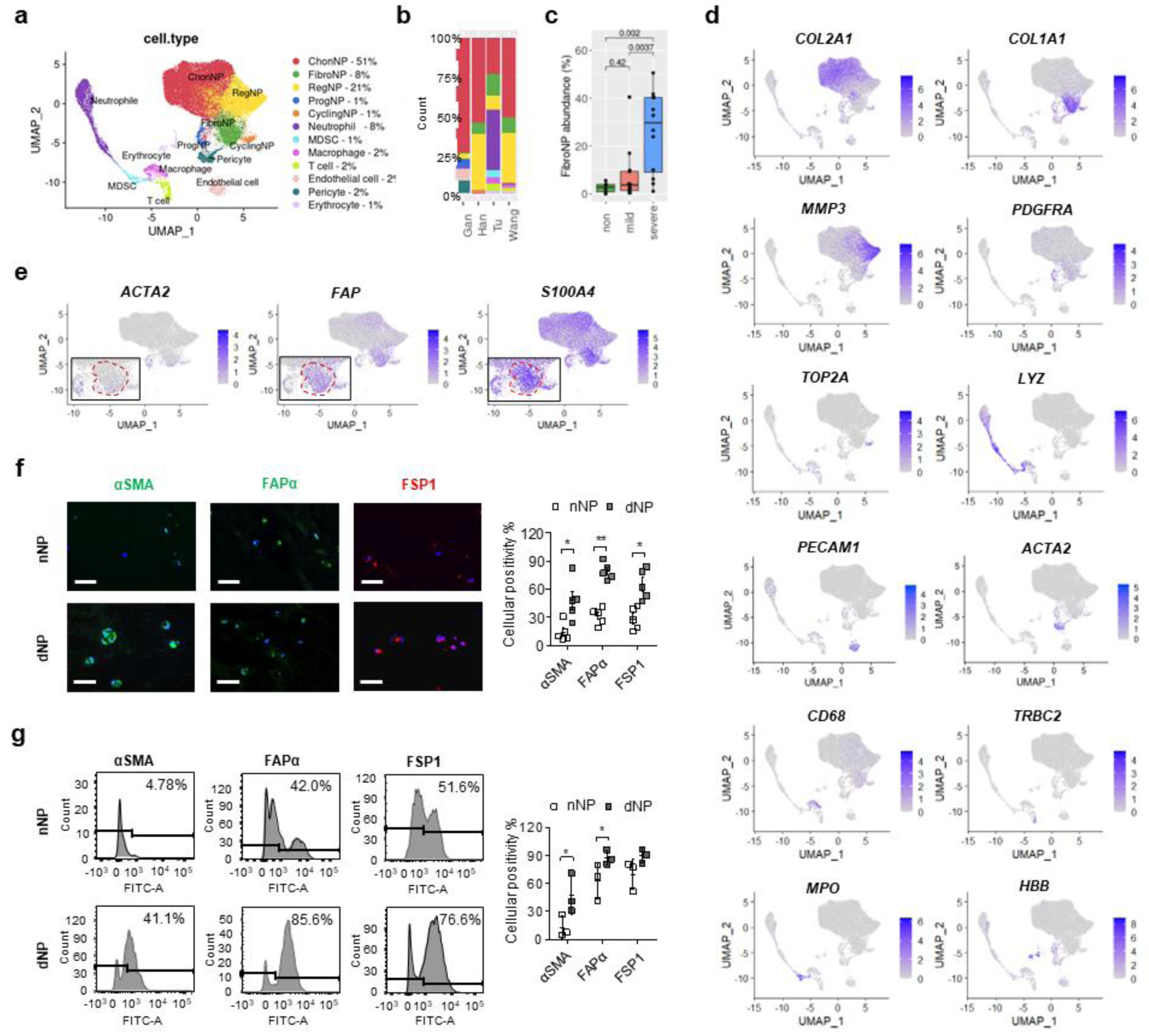
Integrative single-cell transcriptome analysis and identification of fibroblastic NPC in human disc degeneration. **(a)** Identification of 12 distinct cell populations by unsupervised clustering. **(b)** Distribution of the cell populations in individual datasets. **(c)** Degeneration-related FibroNP abundance. **(d)** Signature gene expression for cell clusters defined in A. **(e)** Expression distribution of fibroblastic markers *ACTA2*, *FAP*, and *S100A4*. **(f)** Immunofluorescence and quantification of fibroblastic marker expression in human NP tissues (n=5). Scale bars: 50μm. Nuclei were counterstained with DAPI (Blue). **(g)** Representative flow cytometry plots to identify myofibroblastic NPC. Percentage of αSMA^+^, Fap-α^+^ or Fsp1^+^ cells was averaged from three independent experiments. *p < 0.05, **p < 0.01, and ***p < 0.001 by two-tailed unpaired *t*-tests.

**Table 1.**
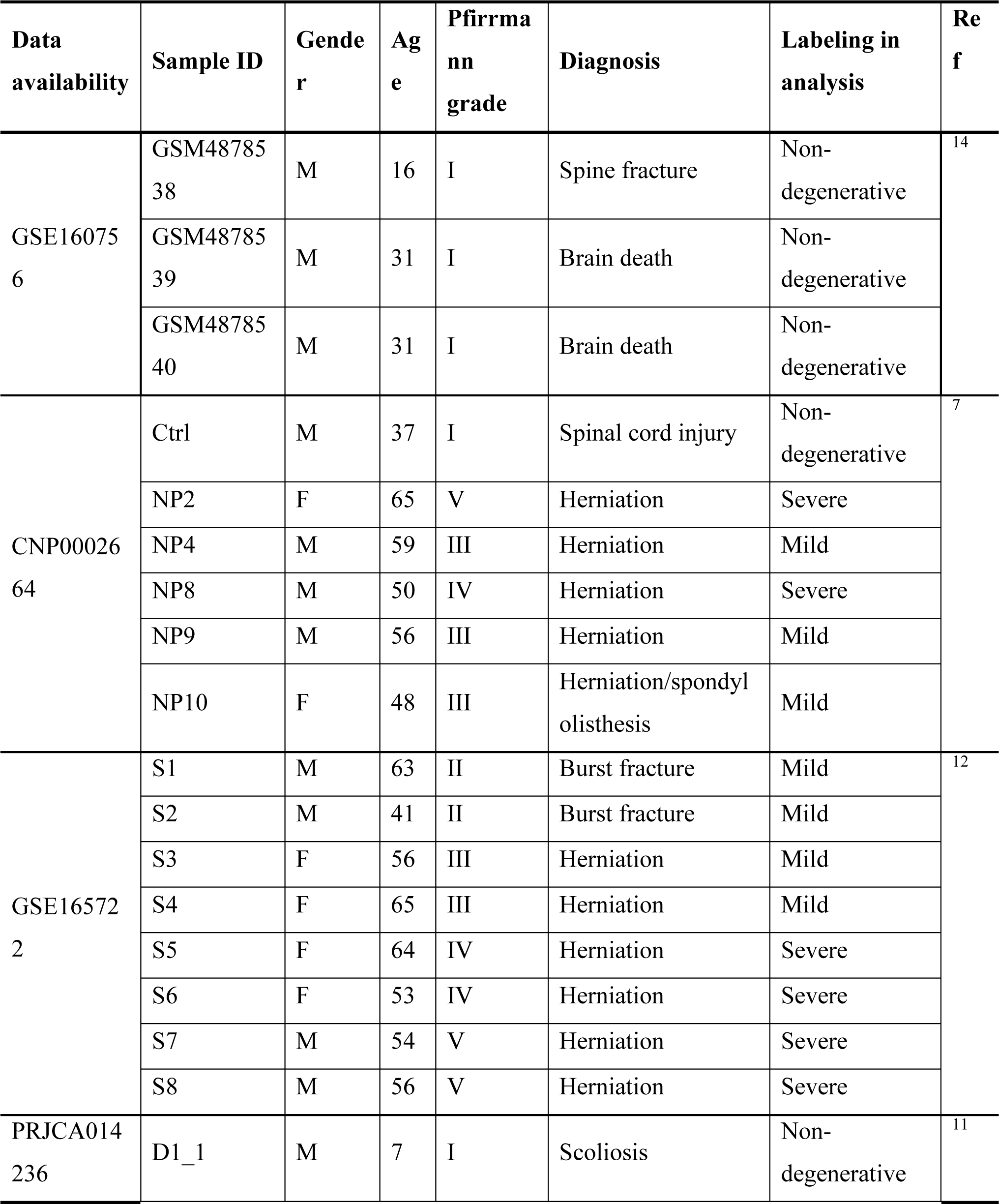
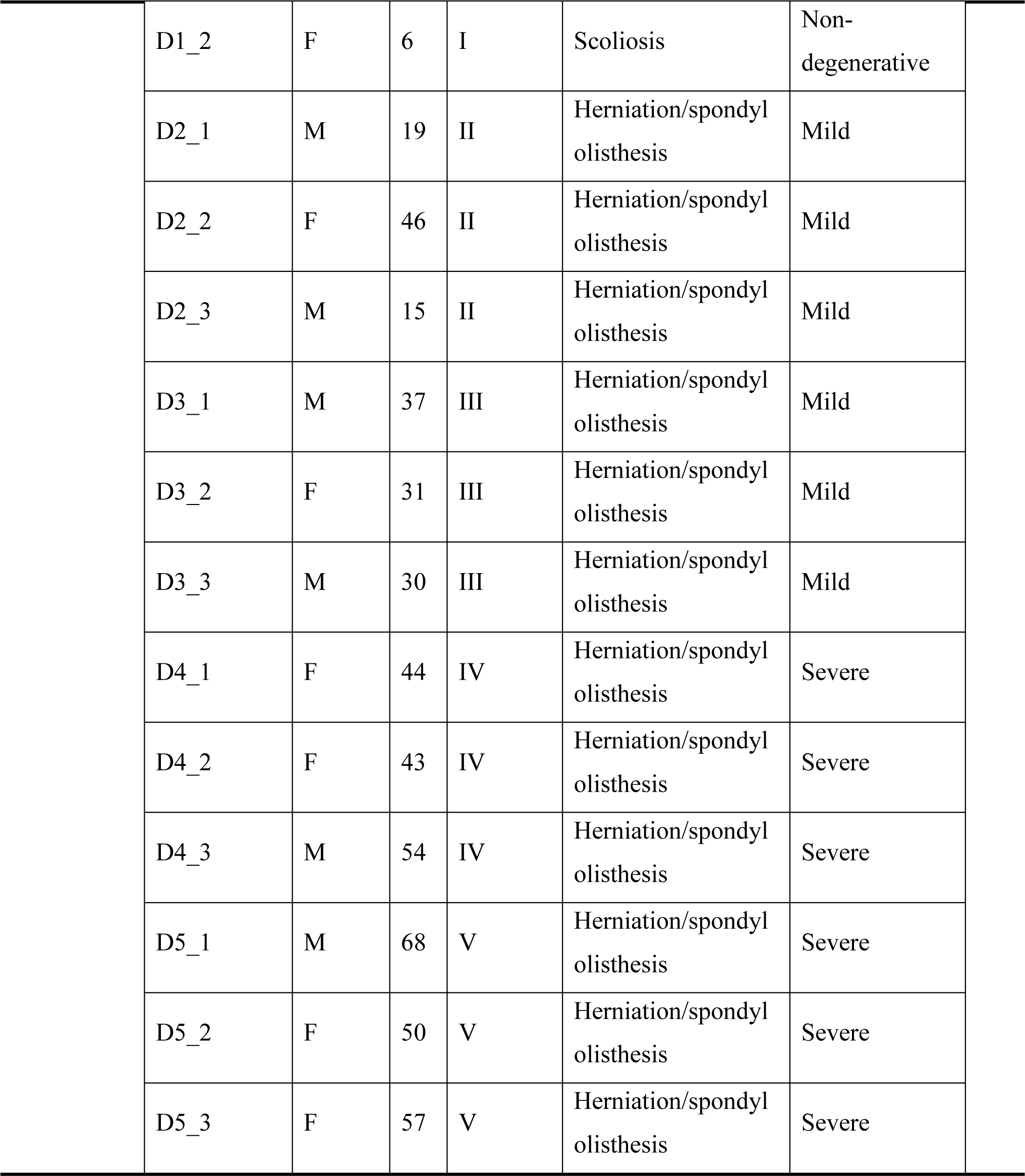
Demographic data. A total of 31 subjects from 4 studies were selected with degeneration graded I-V, and their single cell RNA sequence data were extracted and integrated for further analysis. Samples defined as Pfirrmann grade I were considered as Non-degenerative; grades II/III and IV/V as respectively Mild and Severe degeneration. F: Female; M: Male.

Contractile myofibroblasts are primary effector cells of tissue fibrosis^8^. FAPα and FSP1 mark the active fibroblasts or myofibroblast precursors^17^, and αSMA marks the mature myofibroblasts^18^. These three markers were found enriched in the FibroNP (**Fig. 1e**). More fibrillar collagens, specifically thick collagen I and thin collagen III, were deposited in human degenerative IVDs, which together with reduction of hyaline matrix aggrecan and collagen II, indicated NP fibrosis (**Supplementary Figure S3**). Immunofluorescence showed that the number of cells expressing αSMA (14.0% vs 48%), FAPα (31.4% vs 78.8%) and FSP1 (28.4% vs 65%) in the nNP was far fewer than that in the dNP (**Fig. 1f**), indicating the augmented myofibroblast sprouting in dNP. In line with our single-cell transcriptome analysis, cells from either dNP or nNP tissues were found to contain a lower amount of αSMA^+^ cells than FAPα^+^ or FSP1^+^ cells, which was further supported by fluorescence-activated cell sorter (FACS) analysis of isolated NPC (**Fig. 1g**). FACS also demonstrated an increased myofibroblast population in the dNP compared to the nNP. Notably, nNP (88.2%) and dNP (93.1%) cells presented a comparable amount of cells expressing intracellular collagen I (**Supplementary Figure S4**). However, under monolayer expansion, dNP cells became more flattened and morphologically less refractive, and preferentially expressed higher levels of myofibroblastic markers including *ACTA2*, *FAP* and *COL1A1* (**Supplementary Figure S5**). These findings indicated that myofibroblastic cells became the dominant cell type in the degeneration and supported their contribution to NP fibrosis.

### Disc degeneration contains fibroblastic NPC with hematopoietic features

Previous histological studies suggested that immune cells such as CD68^+^ macrophages and neutrophils may infiltrate into the NP during disc degeneration^19^. In the single cell analysis, immunocyte-like populations could be found at all degeneration grades, expressing markers for neutrophils (*LYZ* and *HLA-DRA*), macrophages (*CD163* and *CD68*), granulocyte-like myeloid derived suppressor cells (G-MDSC) (*ITGAM* and *MPO*) and T cells (*TRAC* and *TRBC2*) (**Table SI**). Interestingly, we found that *COL1A1*, *COL3A1* and *POSTN* were also expressed in the immunocytes, predominantly the neutrophil and macrophage populations (**Fig. 2a**). These *COL1A1+* immunocytes expressed both myeloid lineage genes (e.g. *S100A9*, *PTPRC* and *ITGB1*) and fibroblastic genes (e.g. *VIM*, *TAGLN2* and *COL1A1*; (**Table SI**). Previous studies reported that monocytes contain a subpopulation that co-expresses collagen I and III in addition to markers of hematopoietic cells (CD11b, CD45 or CD34), being referred as fibrocytes^20–22^. The hematopoietic progenitor-associated marker CD34 was immunolocalized with collagen I and contractile protein αSMA in the NP tissue (**Supplementary Figure S6**). However, CD34 is also highly expressed in other cell types such as endothelial cells and muscle satellite cells. *CD34*^+^*COL1A1*^+^ or *CD34*^+^*ACTA2*^+^ cells were in fact mostly enriched in the endothelial cell and pericyte clusters. We further identified *COL1A1*+ subpopulations co-expressing *ITGAM* (coding CD11b) (1094 cells) or *PTPRC* (coding CD45) (1738 cells) (**Fig. 2b**). The abundance of these subpopulations increased with degeneration severity and occupied up to 0.1% cell population dNP tissue (**Fig. 2c**). Immunostaining and flow cytometry verified an increase of CD45^+^COLI^+^ cells in the dNP tissue (16.4% vs 5.4%, **Fig. 2d**) or dNP-derived cell culture (0.80% vs 0.30%, **Fig. 2e**) compared to the nNP controls. These CD45^+^COLI^+^ cells adopted an elongated morphology (insert, **Fig. 2d**). Based on the findings, we regarded the CD45^+^COLI^+^ cells as disc fibrocytes.

**Figure 2.**
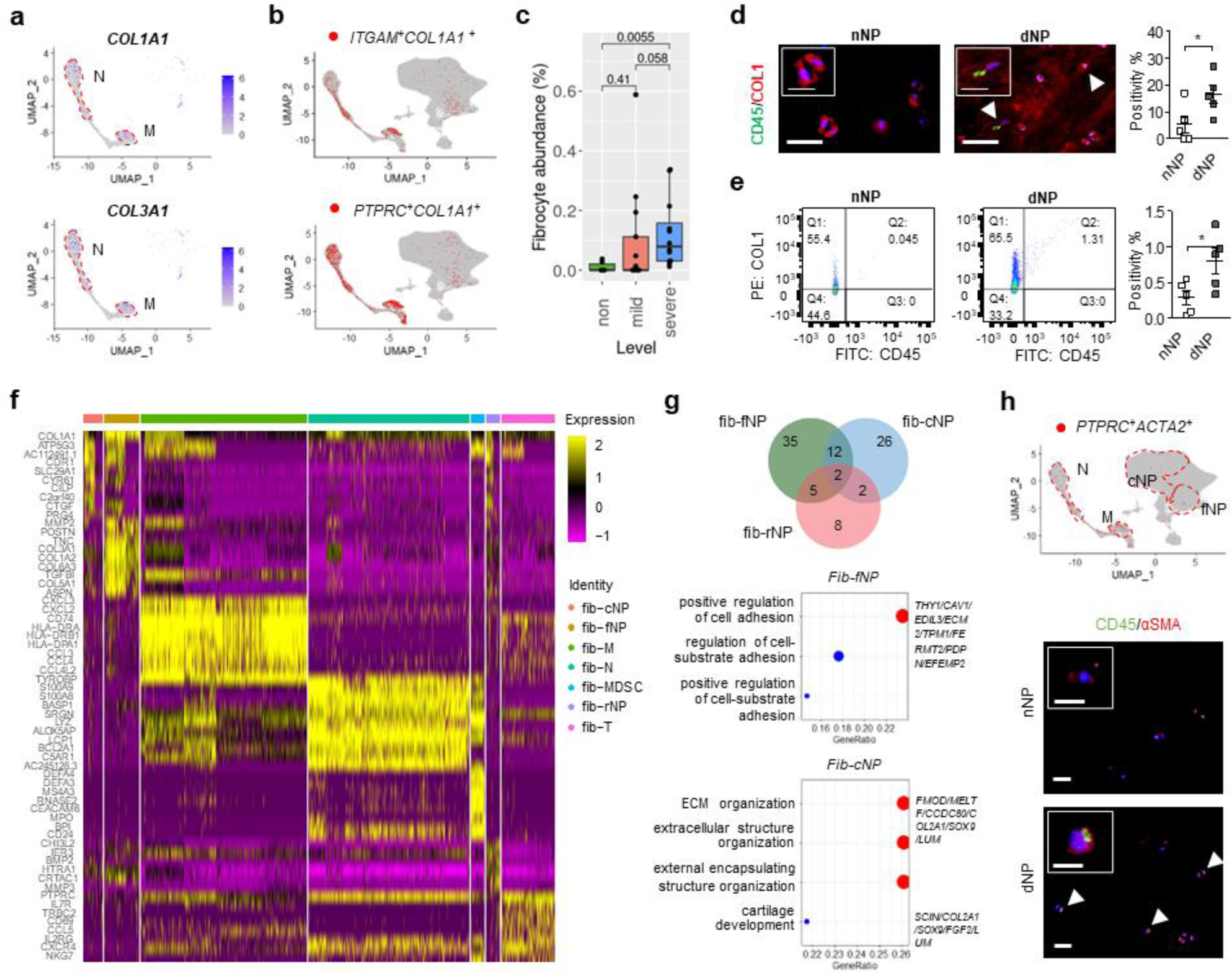
Identification of disc fibrocytes in the disc degeneration. **(a)** Featured gene plots of fibroblast markers of *COL1A1* and *COL3A1* in disc immunocytes. **(b)** Feature plots of expression distribution for monocyte lineage marker *ITGAM* and pan-hematopoietic marker *PTPRC* with *COL1A1*. **(c)** Fibrocyte abundance in disc degeneration. **(d)** Representative co-immunofluorescence micrographs of CD45 and Collagen I (COL1) in human NP tissues from scoliosis (nNP) and degenerative (dNP) IVDs, and quantification of co-expressing cells (n=5). **(e)** Flow cytometric evaluation of disc fibrocytes in isolated human NP cells. Representative flow cytometry was plotted and the percentages of CD45^+^COL1^+^ cells were labeled. **(f)** Heatmap showing and highlighting differentially expressed genes for each disc fibrocytes subtype. (**g**) Venn diagram and GO (gene ontology) functions of fibrocytes residing within three NP cells subclusters: FibroNP (fib-fNP), ChonNP (fib-cNP) and RegNP (fib-rNP). **(h)** UMAP distribution and immunodetection of *PTPRC (coding CD45)*^+^*ACTA2 (coding* αSMA)^+^ cells sin disc NP tissues. Arrowhead: CD45^+^ αSMA**^+^** cell; sale bar: 50 μm. Insert: representative staining micrograph showing positive cellular signals; scale bar: 12μm. *p < 0.05, **p < 0.01 by two-tailed unpaired *t*-tests.

We identified disc fibrocytes within the NPC clusters, present mostly in the FibroNP (7.5%, as fib-fNP), ChonNP (4.2%, as fib-cNP) and RegNP (3.0%, as fib-rNP), in addition to immunocyte clusters of macrophages (35.4%, as fib-M), neutrophil (34.2%, as fib-N), T cell (11.4%, as fib-T) and G-MDSC (3.0%, fib-G) (**Supplementary Table S2**). By comparing these fibrocyte populations, we identified specific gene markers (**Table SI**) and examined their expression (**Figure 2f**). *GNB1* and gluconeogenesis-related gene *ALDOA*, together with marker genes of *COL1A1* and *ITGAM*, were enriched in both immunocytes-(fib-M/N) and NPC-clustered (fib-cNP/fNP) fibrocytes. Within immunocyte-clustered fibrocyte subpopulations (fib-M/N/T/G), a total of 47 genes (e.g. *PTPRC* and *CAPZA1*) were considerably enhanced and mainly linked to the actin organization and immune process (**Table 2 and Supplementary Figure S7**). *RORA* was dominantly expressed in fib-T. Fib-M-related genes *LGALS1* and *LGALS3BP* have been linked to monocyte-derived fibrocyte differentiation^23^. 43 genes (e.g. *COL3A1*, *ASPN* and *ITGB5*) that are commonly expressed in the three NPC-derived fibrocyte subsets are related to the response to TGF-β (**Supplementary Figure S7**). This finding is consistent with the notion that, similar to other fibrotic diseases, disc fibrocyte maturation may involve TGF-β-mediated endothelial-mesenchymal transition^24^. We also performed GO analysis for DEGs specific to the NPC-fibrocytes (**Table 3**). 35 genes distinctly enriched in fib-fNP were mostly related to cell adhesion function. 26 genes enriched in the Fib-cNP were related to matrix organization. Two degeneration-featured genes, *HTRA1*^25^ and *ANGPTL4*^26^ were distinctly expressed in the NPC-fibrocytes (**Fig. 2g**). Fibroinflammatory molecules of TGFBI and SPP1 from the disc fibrocytes were inferred to act on the FibroNP and stimulate the fibrotic process using CellChat analysis (**Supplementary Figure S8**). Fibrocyte-FibroNP communication may also rely on POSTN-ITGB5. Moreover, *ACTA2* was found expressed in *PTPRC*^+^ FibroNP cells (**Fig. 2h**), supporting their potential in myofibroblastic differentiation^23,24^. Consistent with the scRNA-seq finding, co-immunostaining showed a markedly higher expression of CD45 and αSMA in the dNP compared to the nNP (**Fig. 2h**). We also observed αSMA^+^ cells negative for CD45.

**Table 2.**
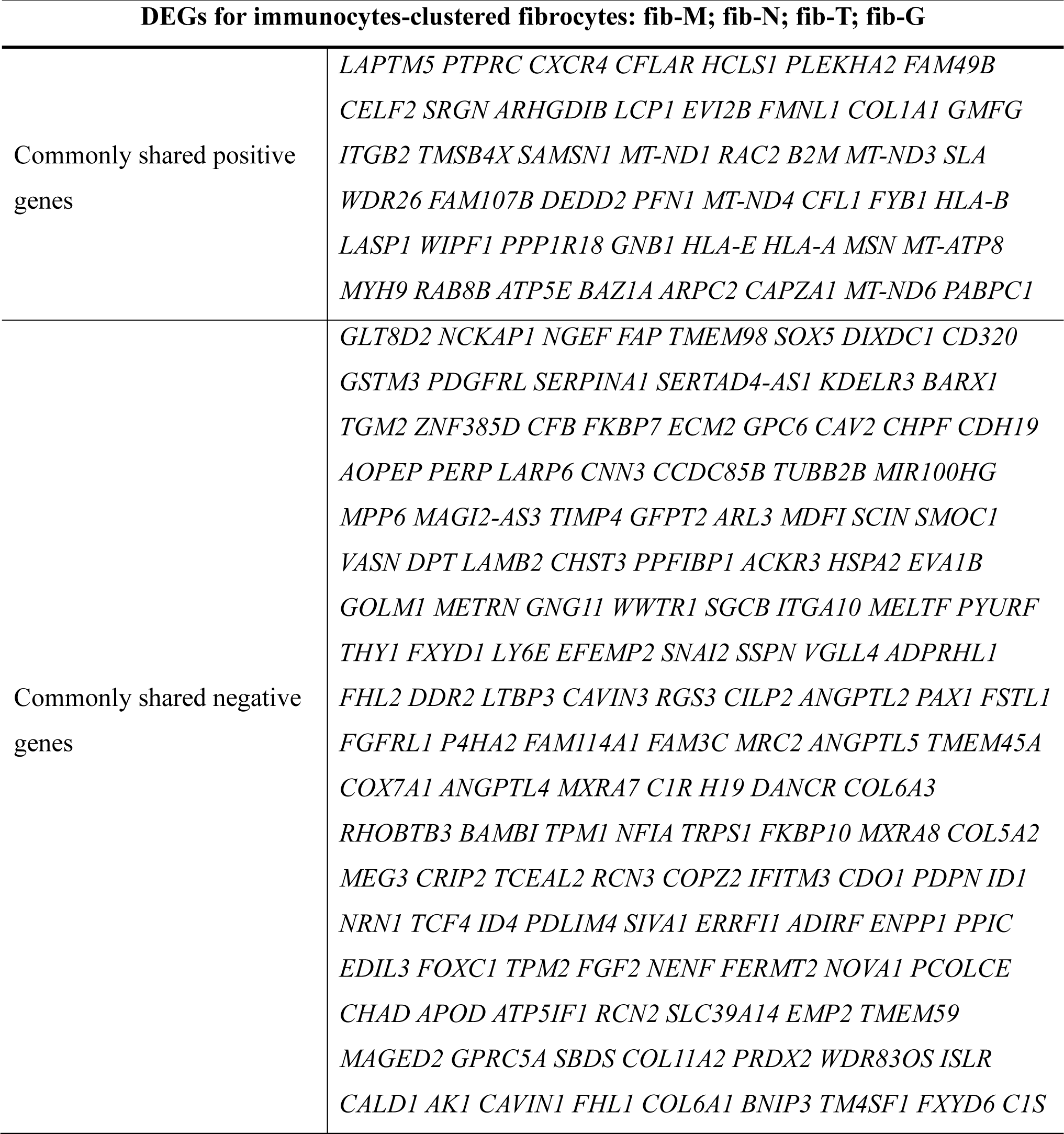
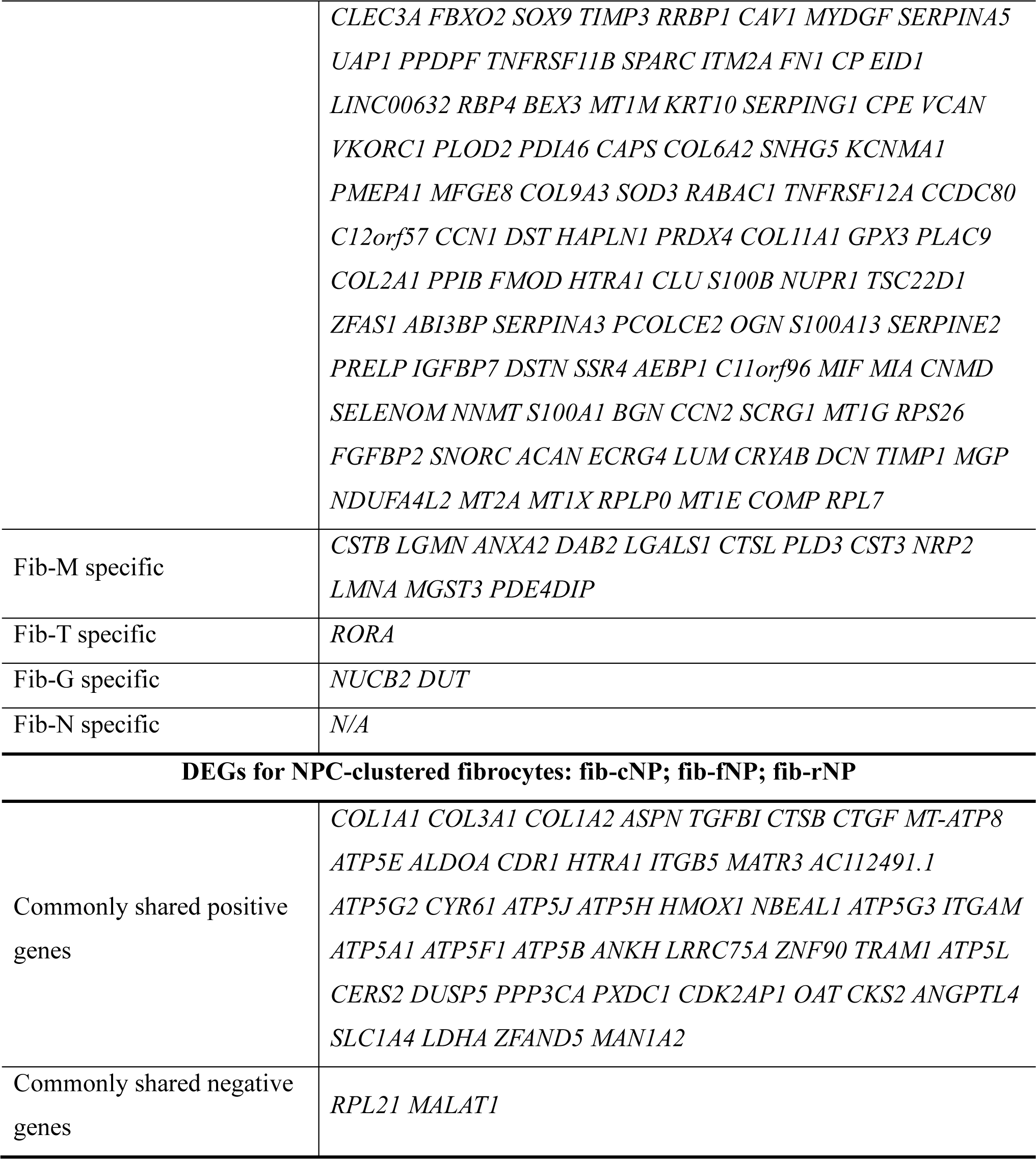
Analysis of DEGs for disc fibrocytes. *PTPRC^+^COL1A1^+^* or *ITGAM^+^COL1A1^+^*NPC were considered as disc fibrocytes, and subdivided to two major clusters based on their UMAP distribution: 1) immunocytes-clustered fibrocytes containing fib-M (Macrophage), fib-N (Neutrophil), fib-T (T cells) and fib-G (G-MDSC); 2) NP cells (NPC)-clustered fibrocytes containing fib-fNP (FibroNP), fib-cNP (ChonNP), fib-rNP (RegNP). Positively and negatively expressed DEGs of different fibrocytes subsets were demonstrated.

**Table 3.**
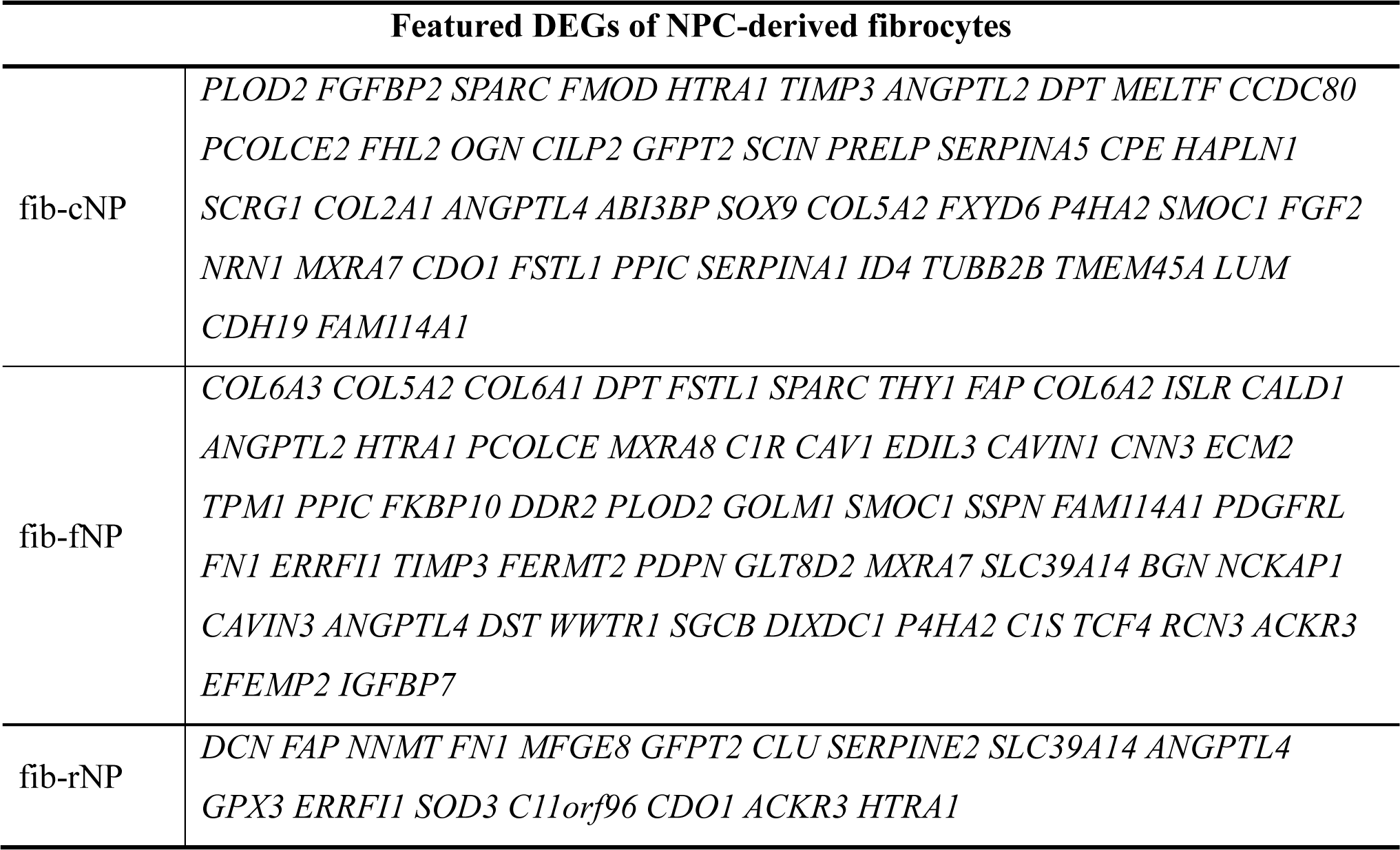
Analysis of DEGs for NPC-clustered disc fibrocytes. NP cells (NPC)-clustered fibrocytes were subdivided based on their UMAP distribution: fib-cNP (ChonNP), fib-fNP (FibroNP) and fib-rNP (RegNP). Featured genes of each subset that were negatively expressed by immunocytes-clustered fibrocytes were summarized.

Given the fibrocytes being a lineage of monocytes expressing CD11b, we studied their temporospatial distribution in the puncture injury induced disc degeneration in CD11b-DTR mice, in which the expression of EGFP-diphtheria toxin (DT) receptor fusion protein is controlled by the *ITGAM* promoter, thereby directing transgene expression and expressing membrane-localized GFP in monocytes/macrophages^22,27^ (**Supplementary Figure S9**). Normal mouse NP, which contains predominantly notochordal cells^4^, was negative for GFP expression (**Supplementary Figure S9**).

Annulus puncture injury led to disc degeneration with NP fibrosis^3,4^. We observed an emergence of GFP^+^ cells in the NP (∼15% of NPC population) by day 3 after puncture, which continued to peak at roughly 35% after 2 weeks and gradually subsided thereafter (**Fig. 3a**). GFP^+^ cells were also observed in the annulus fibrosus (AF) (**Supplementary Figure S9**). We could observe GFP^+^ cells that barely expressed CD11b in the NP at 2 weeks post-puncture (2wpp, **Fig. 3b**), possibly because of a longer turnover of GFP compared to CD11b^27^. Up to 95% of the GFP^+^ NPC were CD45-positive, indicating their myeloid origin (**Fig. 3c**). In line with previous report of fibrocyte differentiation into myofibroblast, we observed αSMA expression in nearly 60% GFP^+^ cells at 4wpp (**Fig. 3d**). The abundance of these dual-positive cells decreased significantly by 8wpp, indicating that the emergence of fibrocytes might be transient or they underwent differentiation. Furthermore, we detected CD45^+^COLI^+^ and CD45^+^αSMA^+^ cells in the punctured disc (**Fig. 4b&c**), and the amount of CD45^+^αSMA^+^ NPC showed a two-fold reduction by 8wpp compared to 4wpp. These findings collectively support the notion of fibrocyte infiltration into the degenerating discs and adopting a myofibroblastic phenotype (CD45^+^αSMA^+^) during NP fibrosis.

**Figure 3.**
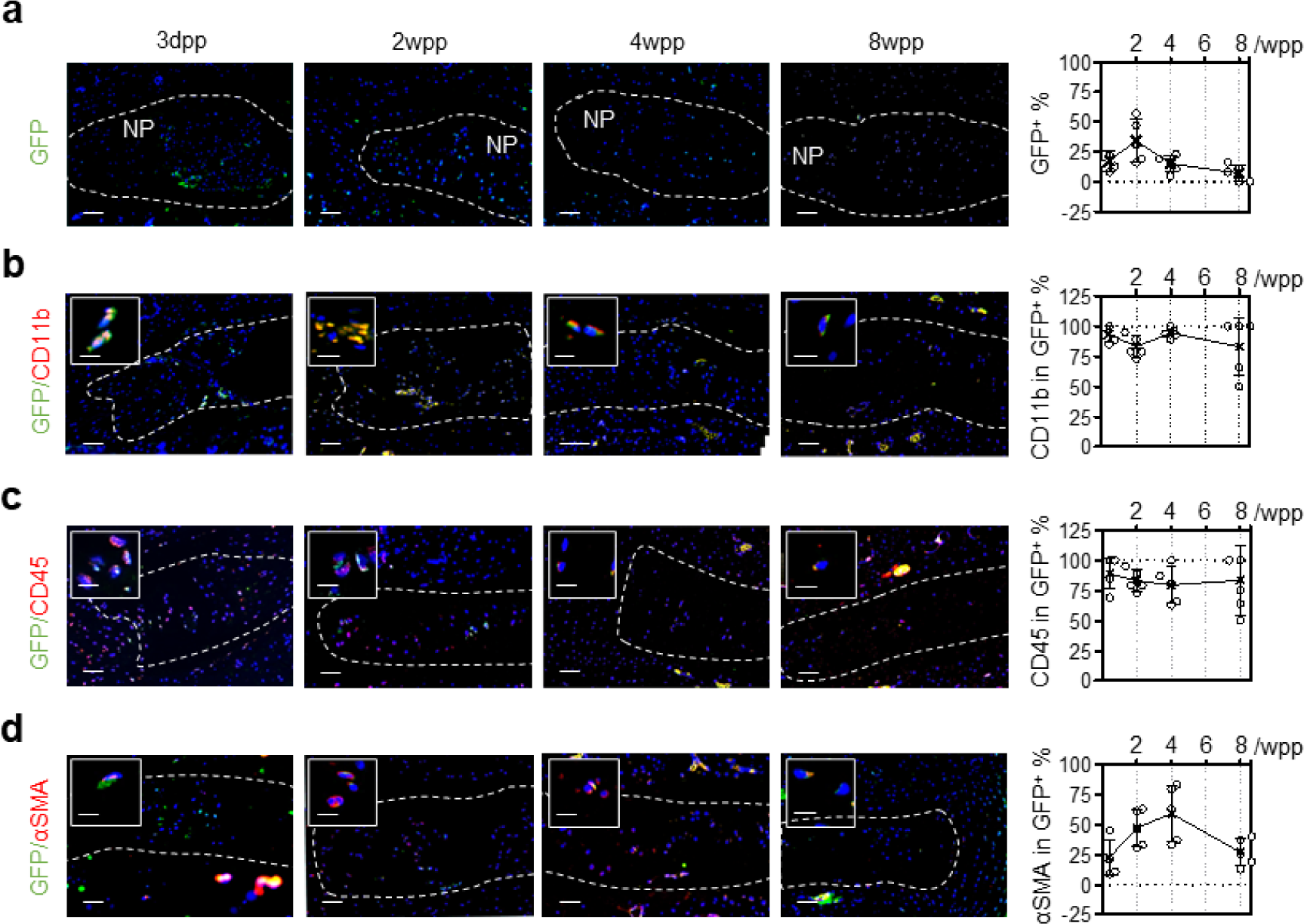
Hematopoietic fibrocytes contribute to myofibroblastic NPCs. **(a)** Immunostaining and quantification of GFP expression in the degenerative caudal discs of CD11b-DTR in 3 days (dpp) to 8 weeks post-puncture (wpp) window. **(b)** Co-immunostaining of hematopoietic marker CD45 and **(c)** αSMA in CD11b^+^ cells with GFP with quantification (GFP^+^CD34^+^, **d**; GFP^+^αSMA^+^, **e**). Cellular positivity counting was conducted in five representative sections, and expressed as the mean ± S.D. Sale bar: 50 μm. * p<0.05, One-way ANOVA with Bonferroni post-hoc test.

**Figure 4.**
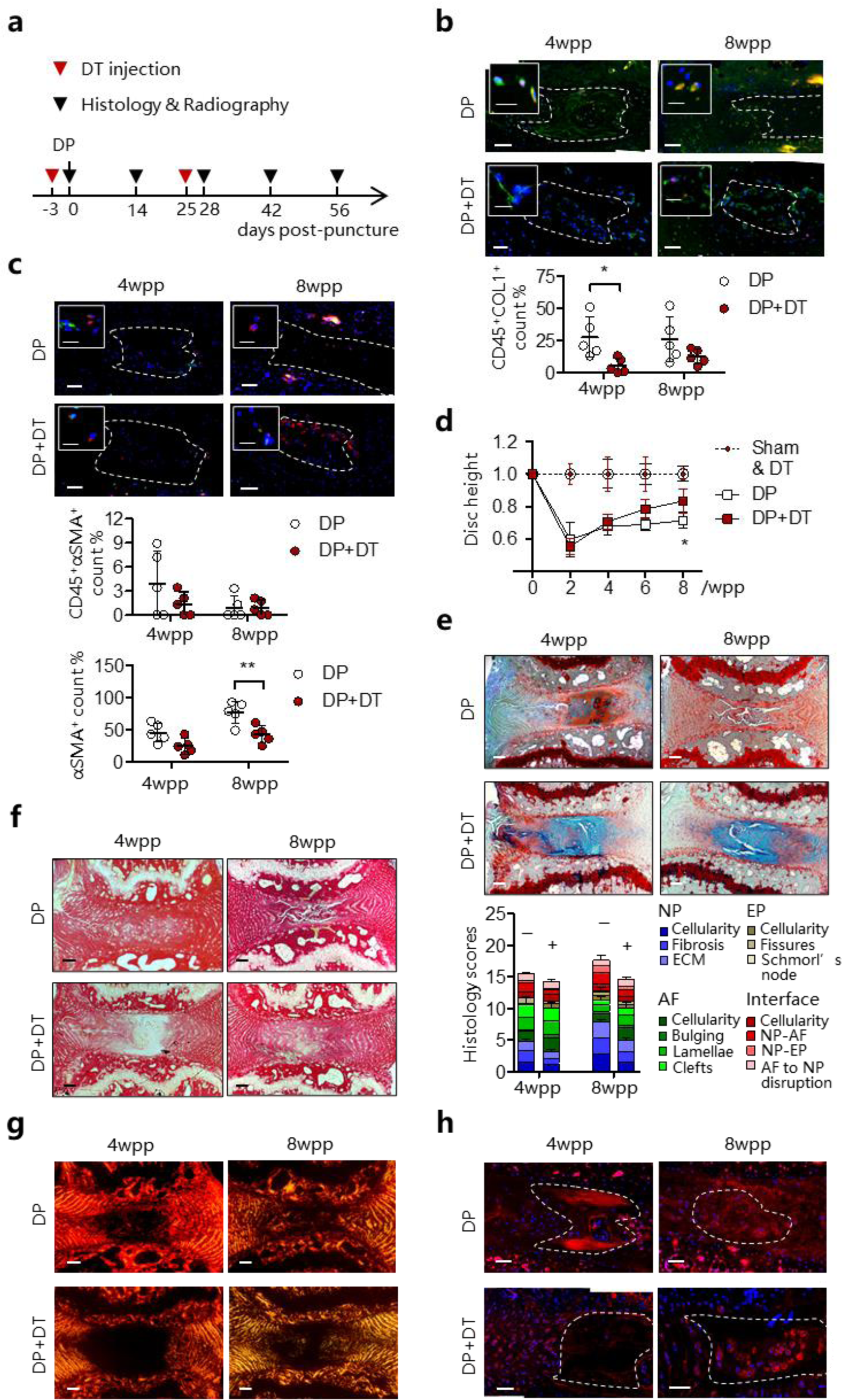
Fibrocyte deficiency suppresses disc degeneration and NP fibrosis. **(a)** Time-line illustration for DT administration and puncture surgery on CD11b-DTR mice. PBS serves as vehicle control. **(b)** Co-staining of CD45 (red) with COL1 (green) or αSMA **(c,** green) and dual-positive cells counts. αSMA positivity in NPC was also counted. **(d)** Time course assessment of disc height changes (n=5 per group). **(e)** FAST staining and degeneration scoring based on compartmental deformation in NP, annulus fibrosus (AF), endplate (EP) and interface. **(f)** Picrosirius red staining and **(g)** their polarized microscopy. **(h)** Collagen I immunostaining. In immunofluorescence, nuclei were counterstained by DAPI. Scale bar: 50μm, insert scale bar: 12μm. Dot line: NP region. Wpp: weeks post-puncture; DP: disc puncture. Data are expressed as mean ± S.D. Two-way ANOVA with Bonferroni post-hoc test, * p<0.05.

### Fibrocyte depletion rescues NP fibrosis

We tested whether depletion of CD11b-expressing cells by DT administration^27^ could lead to a reduction of fibrocytes in the puncture induced IDD. We first examined the frequency and dosage of DT injection to enable extended depletion while minimizing lethality due to compromised immunity (**Supplementary Table S3**). The number of CD11b^+^ monocytes were largely reduced in peripheral blood (>50%) after twice administration of DT at 10μg/kg (**Supplementary Figure S10**). For the IDD model, the first injection was performed 3 days prior to the puncture surgery (**Fig. 4a**). We found a marked decrease of the GFP^+^ cells in the NP at 2 and 8wpp, supporting the effect of DT on the transgene expressing cells (**Supplementary Figure S10**). Interestingly, a lower number of CD45^+^COL1^+^ cells was observed in the NP after the second dose of DT injection by 4wpp when compared to vehicle control (**Fig. 4b**). The amount of CD45^+^αSMA^+^ NPC also tended to decrease although the difference is not statistically significant (**Fig. 4c**). These findings indicate that DT administration caused a depletion of monocytic fibrocytes in the NP which emerged during injury-induced IDD.

We questioned if the loss of the myeloid derived fibrocytes leads to a modification of IDD and NP fibrosis. Puncture induced IDD showed a significant disc height loss by 50% at 2wpp and 30% at 8wpp (**Fig. 4d**). DT administration could induce a gradual recovery and reduce the disc height loss by >80% at 8wpp. Disc integrity assessed by FAST staining^3^ showed a prominent improvement with DT administration at both 4 and 8wpp, including reduced lacune formation, preserved NP-AF boundary, and increased alcian blue intensity in the NP (**Fig. 4e**). Moreover, alcian blue staining in the NP of the DT-injected mice was largely maintained, implying that the loss of proteoglycans was prevented. The amount of myofibroblastic NPC positive for αSMA steadily increased after puncture and took up to 76% of the whole population by 8wpp (**Fig. 4c**). DT administration could lower its positivity to 41%. It is noteworthy that the majority of αSMA^+^ cells detected in the punctured discs, in particular in the DT-treated group, was negative for CD45 (>90%). Picrosirius red staining and immunostaining of collagen I further consolidated a decrease in fibrillar collagens (**Fig. 4f-h**).

Pericellular expression of collagen I was noted in the NP after DT administration. Collectively, these findings demonstrated that IDD progression is dependent on a function of myeloid derived cells where the CD45^+^COL1A1^+^ fibrocytes and αSMA^+^ myofibroblastic cells may mediate NP fibrosis.

## Discussion

NPC are conventionally defined as collagen II-, aggrecan- and Sox9-expressing cells. Recent studies of single-cell transcriptome have indicated cell heterogeneity in the human NP, which contains cell populations expressing notochordal, chondrogenic, and fibroblastic phenotypes^7,11–14^. Our meta-analysis of the human disc scRNA-seq datasets has outlined the commonly featured NPC hierarchy and stratified their fibroblastic subtypes. *COL1A1*^+^ FibroNP is previously defined as adhesion/fibroNPC^12^ or C4 NPC^7^. The FibroNP population was not reported in Wang et al scRNA-seq dataset, possibly because their cells were derived from the whole disc tissue without NP isolation and therefore may have dominated by *COL1A1^+^* AF cells^11^. Nonetheless, they defined the FibroNP genes (*POSTN*, *COL1A1*, *COL3A1* and *TMSB4X*) in MK167^+^ progenitor and NP progenitor cells (**Supplementary Table 1**), implying the fibroblast presence within the two NP subtypes. Han et al found a small fraction of fibrochondrocyte progenitors in the NP, which partially constitute the aforementioned two NP subsets. The regulatory NPC and C1/3 chonNPC subtypes reported by Tu^12^ and Han^7^ are merged to the RegNP cluster in this study, sharing the expression of *MMP3*, *CHI3L2*/*1* and *GPX3*. The CyclingNP cluster is a pool of *TOP*2A^+^ cells fraction from the *COL2A1^+^* ChonNP or *COL1A1^+^* FibroNP subsets, implying they may be derived from a common replicative progenitor origin. Pseudotime ordering of the NP clusters and velocity analysis may help construct their relationship.

The combination of collagen production (e.g. collagen I) and the expression of leukocytic common antigen (CD45) or one of the hematopoietic or myeloid antigens (e.g. CD34 or CD11b) is considered as a sufficient criterion for fibrocyte identity^21^. *CD34^+^COL1A1*^+^ cells are mainly enriched in the endothelial cells and pericytes subpopulations (**Supplementary Figure S6**). *ITGAM* (coding CD11b) or *PTPRC* (coding CD45) is therefore preferrable to CD34 as the marker of fibrocytes in human NP tissues. Our single cell transcriptome analysis showed a wider UMAP distribution and higher number of *PTPRC^+^*than *ITGAM^+^* cells, supporting the notion that CD11b is a more selective yet less representative fibrocyte marker compared to the pan-hematopoietic markers Vav-1 or CD45^21^.

Pronase digestion was commonly deployed for releasing single cells from the NP in scRNA-seq and flow cytometry assays^28^. However, the digestion can also results in extensively membrane proteins degradation, and thereby affect the antibody recognition of surface molecules^29^. This may in part explain the discrepancy in measurements of CD45^+^COL1A1^+^ cells between in situ immunohistochemistry and flow cytometry (**Fig. 2d&e**).

In addition to the NP clusters, *CD11b^+^COL1A1*^+^ or *CD45^+^COL1A1*^+^ cells are mostly located within the macrophages, neutrophils and T cells clusters, implying the fibrocytes may be derived from multiple sources or differentiation routes of the immune system. Blood immune cells are casually linked with IDD, such as macrophages^19^ and CD39^+^CD4^+^ T cells^30^. Interestingly, differentiation of monocyte (CD11b^+^CD115^+^Gr1^+^) into fibrocyte could be dependent on CD4^+^ T cells activation^31^.

Immune cells can infiltrate through focal defects in cartilaginous endplate (EP) and AF advancing the fibroinflammatory process^32^. Monocytes are defined as *CD45^+^CD11b^+^*cells and constitute the myeloid-derived macrophages (*CD45^+^CD11b^+^F4/80^+^*), neutrophils (*CD45^+^CD11b^+^HLA-DRA^+^*) and MDSCs (*CD45^+^CD11b^+^OLR1^+^*)^12,13,21^. G-MDSCs, shown in cardiac fibrosis with myofibroblast activation capabilities, contribute to T-cell suppression and ROS production in IDD^12^. Monocyte-derived inflammatory macrophages could transit into myofibroblasts^33,34^ and were identified in the degenerative NP^19,35^. CD11b^+^CD45^+^ monocytes were previously reported to support the proliferation and colony forming ability of in vitro-cultivated MSCs^36^. Whether the monocytic cells infiltrated into the degenerating discs may exert a similar effect on local mesenchymal progenitors to promote fibrogenesis or repair is not clear. Moreover, CCR2^+^ monocytes have been identified in the degenerative IVDs from herniation patients and constitute the majority of CD11b^+^ and F4/80^+^ cells^37^. However, as reported in myocardial infarction model^38^, loss of CCR2 expression is required for fibrocyte differentiation of the infiltrated monocytes. This is in line with our observation that the disc fibrocytes are negative for *CCR2*. In a mouse model of renal fibrosis where CCR2 is depleted, migration of circulating fibrocytes to the kidney was interfered^27^. Whether CCR2 plays a role in monocyte infiltration and hence differentiation into disc fibrocytes await to be determined.

Fibrocytes could be derived not only from local monocyte differentiation but also migrated from peripheral blood in the form of partially differentiated collagen-producing cells^21,27^. Recruitment of monocytes and peripheral blood fibrocytes may rely on disc neo-vascularization. Normally, blood vessels recede from the outer AF and the EP in adults. However, under degenerative conditions the NP can become re-vascularized and blood vessel infiltration is frequently observed and spatially associated with immunocytes infiltration^39^. Most of the degenerative NP tissues in this study for scRNA-seq analysis were dissected from extruded NP of herniated discs that normally show pronounced neo-vascularization^40^. The increased number of GFP^+^ cells found in the AF and EP of the punctured CD11b-DTR discs may indicate the infiltration through the AF and EP routes.

Pericytes (CD34^+^αSMA*^+^*) were immunodetected in the degenerative human NP (**Supplementary Figure S6)**, supporting the presence of neo-vascularization. In fact, pericytes can contribute to scar-forming myofibroblasts in kidney and subretinal fibrosis^8^. Studying disc vascularization and its relation to the emergence of disc fibrocytes may provide further insights into the pathway of NP fibrosis.

Trafficking fibrocytes express chemokine receptors CCR5^41^, CCR7^21^ and CXCR4^21,42^. Studies have reported increased CCR5 expression in AF and NP cells^43^ and chemokines CCL2/7 in degenerated NP^44^. Bone marrow cells, such as CD146^+^ MSCs^45^, may have a potential to migrate into the NP, possibly by control of SDF1^46^. CCL5 could interact with CCR1/3/5 in NPC and was up-regulated under inflammatory stimuli^47^. CXCR4 is a receptor of SDF1^48^ and CXCL12^42^, which was found upregulated in degenerated discs and implicated in disc angiogenesis^48^ and cellular apoptosis^49^ via PI3K/NF-κB pathway. Interestingly, we find that the disc fibrocytes highly express *CXCR4* and *CCR1* (**Table 2**), and may communicate with the ProgNP population via the CXCL12-CXCR4 axis. Future study may test if CXCL12 has a function to recruit *CD45^+^COL1A1^+^*fibrocytes.

Tissue fibrosis is mediated by myofibroblast from local or extrinsic sources^8^ and fibrocytes may serve as one of the progenitors of myofibroblasts^21,24^. Expression of αSMA is the hallmark of mature myofibroblasts and essential to their cellular contractility^18^. Expression of αSMA in NPC may therefore indicate their myofibroblastic property^24^. FAP-α is a well-known biomarker of (myo-)fibroblasts, and together with FSP1 it can label most of the collagen I-producing cells in the bone^50^. FSP1 marks an active fibroblast population and represents a non-overlapping fibroblast entity to αSMA^+^ subtype in heart, kidney and skin fibrosis disease^51^. FSP1 is also expressive in atypical myeloid-derived F4/80-positive inflammatory macrophages in liver injury^52^. FSP1 was found expressed in the *COL2A1*^+^ ChonNP cells in addition to the *COL1A1*^+^ FibroNP cells.

Our results demonstrate αSMA expression in the CD45^+^ and CD11b^+^ fibrocytes, supporting the conversion of fibrocytes to myofibroblasts in NP fibrosis. Yet, a vast number of αSMA^+^ myofibroblastic NPC in both human degenerative IVDs and mouse punctured discs are not positive for CD45 and CD11b. In the CD11b-DTR mice where the monocytes and their derived lineages (including fibrocytes) were depleted^27^, αSMA expressing cells could still be observed (>55%, 2 wpp in **Fig. 4d**) in the degenerative NP. This may be due to the existence of non-hematopoietic origins of the myofibroblast pool, such as those derived from resident NPC^4^. Alternatively, fibrocytes may lose CD45 and CD11b expression during fibrocyte differentiation^33^. We also note the presence of residual CD45^+^COL1^+^ cells and GFP^+^ cells after DT administration. This likely arises from the incomplete monocyte ablation and a continuation of fibrocyte recruitment.

Fibrocyte-to-myofibroblast conversion acquires activation of TGF-β/SMAD signaling^53^, and interleukin-18 receptor 1^54^ and muscarinic receptor M3^55^ are shown to promote the fibrocyte formation and contractile function. Moreover, fibrocytes can secret soluble factors such as periostin^56^ and TIMP1^57^ and semaphorin-7A^58^ to regulate myofibroblast activity and collagen expression in pulmonary and intestinal fibrosis^20^. Class 3 semaphorins expression are found associated with disc innervation and angiogenesis^59^. Interestingly, TIMP1^60^ and POSTN^61^ are up-regulated in the degenerated discs and enriched in the FibroNP. Our CellChat data suggest that POSTN may interact with ITGB5 of disc fibrocytes. Whether disc fibrocytes regulate NP fibrosis through a paracrine manner awaits to be addressed. Furthermore, the NPC-enriched disc fibrocytes express *HTRA1* and *ANGPTL4*. HTRA1 is a proteolytic enzyme for degrading the matrix either on its own^62^ or by upregulating ADAMTS-5^25^ or MMPs, whereas expression of ANGPTL4 is positively associated with IDD severity ^26^. Notably, our data suggest that ANGPTL4 is a major mediator of the intercellular communication between the disc fibrocytes and the CyclingNP, RegNP and FibroNP subpopulations. Their regulatory role in the fibrocytes and NP fibrosis worths to be investigated.

In conclusion, this study implicates a function of monocytic fibrocytes in NP fibrosis and IDD progression, shedding a new insight to cell fate control of disc immunocytes and their modulatory effect on IVD degeneration. Understanding the effects and regulation of the fibrocytes may potentiate the development of new diagnostic and therapeutic strategies for modifying IDD.

## Materials and methods

### scRNA-seq data acquisition and downstream analysis

Four single cell RNA sequencing data sets (GSE160756, GSE165722, CNP0002664, PRJCA014236) were downloaded and processed for next-step analysis. 31 individual samples were rearranged according to Pfirrmann grade, which Pfirrmann grade I samples were classified as non-degenerative NP, Pfirrmann grade II/III as mildly degenerated NP, and Pfirrmann grade IV/V as severely degenerated NP (**Table 1**). Seurat package (version: 3.1.4, https://satijalab.org/seurat/) was used for cell normalization and regression according to the UMI counts of each sample and percent of mitochondria rate to obtain the scaled data. MNN (mutual nearest neighbor) was used to remove potential batch effect. The top 10 principles were used for unsupervised UMAP construction via graph-based cluster method (resolution = 0.8), and marker genes were determined with p-value < 0.01 and log(fold-change) > 0.25 by the FindAllMarkers function. CellChat analysis was conducted to identify potential intercellular interactions, and probability values for each interaction were calculated. Significant mean and cell communication significance (p-value < 0.01) were calculated based on the interaction and the normalized cell-matrix achieved by Seurat normalization.

### Human samples

All procedures described in the present study were approved by the Institutional Ethics Review Board of The University of Hong Kong. IVD tissues were obtained from patients under informed consent. Non-degenerative (n) and degenerative (d) IVDs were harvested from respectively adolescent idiopathic scoliosis (n=7) and lumbar disc degeneration patients (n=9). The age of the patients ranged from 14 to 72 years, and tissue was evaluated with the Pfirrmann classification system (**Supplementary Table 4**).

### Cell isolation and culture

Jelly-like NP tissues were carefully dissected from the cartilaginous EP and lamellar-structured AF. In severely degenerated IVDs (grade IV-V), NP tissues were more fibrotic and hardly distinguishable from inner AF, so we harvested the tissue presumably at the central region of the NP. Primary NPC were extracted by sequential enzymes digestion of pronase (0.25%, Roche Diagnostics) and collagenase II (600U/ml, Worthington Biochemical)^28^, and maintained in high-glucose Dulbecco’s modified Eagle’s medium (DMEM) (Invitrogen) supplemented with 10% fetal bovine serum (FBS, Biosera) and 1% penicillin/streptavidin (P/S, Invitrogen) at 37 °C and 5% CO_2_.

### Flow cytometry

Newly extracted NPC were washed by PBS and re-suspended in PBS. Cells were fixed in 4% paraformaldehyde (PFA) and blocked by 5% BSA. Permeabilization with 0.1% triton X-100 (Sigma-Aldrich) was applied where necessary. Cells were then treated with primary and secondary antibodies, and signals from cells labeled with conjugated fluorophores were detected by using BD FACS CantoII Analyzer (BD Biosciences). The antibodies used for different flow cytometry analysis are listed in **Supplementary Table S5**. Appropriate IgG control fluorescence compensation was applied to avoid false positive signals. Data were further analyzed by BD FACS Diva software (BD Bioscience).

### Real-time quantitative PCR

Total RNA was prepared using RNeasy mini kit (Invitrogen) and cDNA synthesis was performed using High-capacity RNA-to-cDNA kit (Applied Biosystems). PCR was performed using PowerUp SYBR green master mix (Applied Biosystems), and data are presented as expression levels relative to *GAPDH* using the 2^−ΔCT^ method. The primer sequences were listed in **Supplementary Table S6**.

### Mouse disc puncture model

Tg (ITGAM-DTR/eGFP)34Lan/J mice express membrane localized GFP fluorescence with diphtheria toxin (DT) receptor under the control of CD11b and were used for disc puncture. The animal experiments were approved by The University of Hong Kong Committee on the Use of Live Animals in Teaching and Research (CULATR). CD11b-DTR mice (female, 12 weeks of age, *n*=5 per group and time points) received needle puncture at levels of C5/6 and C7/8 of tail IVDs as described previously^4^. At four days prior to the timepoints and every other week after the puncture surgery, these mice were subjected to intraperitoneal injection of saline or DT (25μg/kg body weight) (Sigma-Aldrich). At assigned end points, mice were euthanized by intravenous injection of sodium pentobarbital (1.2 g/kg), and intact discs with attached vertebral bones were harvested for histochemistry. Peripheral bloods were harvested at days 0, 3, and 7 after DT injection, and number of GFP^+^ monocytes was measured by fluorescence-activated cell sorting (FACS).

### Disc height measurement

Anterior-to-posterior radiographs of tail discs of CD11b-DTR mice or C57BL/6J were taken using a digital radiography machine (Siemens) with an exposure at 25 kV, for 5 seconds. Disc height index was calculated as previously described ^4^ and presented as percent relative to the unoperated level of tail disc C6/7. Images were analyzed biweekly in all groups up to 8 weeks.

### Histology and immunostaining

Samples from human and CD11b-DTR mice tissues were fixed with 4% paraformaldehyde in phosphate buffer for 24 h. Decalcification was performed for mouse IVDs using mouse solution. Paraffin-embedded tissue blocks of human NP and mice IVDs were sectioned at 7 μm. For histological staining, sections were deparaffinized and rehydrated, and subjected to FAST staining (Alcian blue, Safranin O, Fast green and Tartrazine stain) (Sigma-Aldrich) as previously described^63^, or Masson’s trichrome stain and Picrosirius red stain following standard procedures. Polarized microscopy was combined with picrosirius red staining to observe the collagens network in human NP tissue and mouse IVDs^64^. According to the histomophological changes, disc degeneration in CD11b-DTR mice was scored with reference to Tam’s scoring system^63^.

For immunofluorescence, antigen retrieval was performed following rehydration by incubating with 0.8% hyaluronidase (Sigma-Aldrich) at 37°C for 30 min and antigen retrieval buffer (100 mM Tris, 5% EDTA, pH 9.5) at 95°C for 10 min. The sections were blocked in protein block solution (DAKO) for 60 min before incubating with antibodies. Fluorophore conjugated antibodies were used to enable fluorescence detection. Information for these primary and secondary antibodies were summarized in **Supplementary Table S5**. The sections were mounted with VECTASHIELD Mounting Medium with DAPI for nuclei staining. Isotype-matched mouse, rabbit or rat IgG was used as negative controls. Tissue sections were imaged by fluorescence microscope (Nikon Eclipse E600), and fluorescent positive cells in five fields of view (FOV) were manually counted in ImageJ with reference to total number of cells.

### Data analysis

Data are presented as mean ± S.D. of four to six experiments, as indicated in respective figure legends. Unpaired t-test (two tailed) was used to determine significant differences. Comparisons among multiple groups were assessed using one-way ANOVA with Bonferroni post-hoc test. A p value < 0.05 was considered statistically significant.

## Acknowledgements

This work was jointly supported by the General Research Fund (17121619) of the Research Grant Council of Hong Kong and the Scientific Research Foundation of PEKING UNIVERSITY SHENZHEN HOSPITAL (KYQD202100X).

## Author contributions

YS, YP and ZS contributed equally to this manuscript. YS contributed to the study design, data collection and interpretation, producing figures and obtaining the fund. ZS, KKS and YS conducted the bioinformatic analysis. YS, YP, QL and ML performed the experiments and helped in data analysis. YS, YP, ZS and VYL wrote the manuscript. JZ, YH, ZG, KMC, ZZ and XZ collected samples and clinical data, and edited the manuscript. KMC, ZZ and XZ reviewed the data and provided critical data interpretation. VYL conceptualized and supervised the whole project, provided supports and coordinated the collaboration. All authors have read and approved the manuscript.

