## Supplementary Figures and Tables for "Monocytic fibrocyte-like cell enrichment and myofibroblastic adaptation causes nucleus pulposus fibrosis and associates with disc degeneration severity"

**Supplementary materials**

**Supplementary Figure S1. Signature gene expression for disc pericytes (*RGS5*) and endothelial cells (*CD34*).**

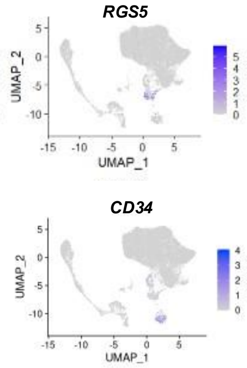


**Supplementary Figure S2. Association of NPC clusters with disc degeneration severity. (A)** Distribution of the five NPC clusters in different disc degeneration grades. **(B)** Abundance of the FibroNP population in different grades. P values were calculated based on unpaired t test.

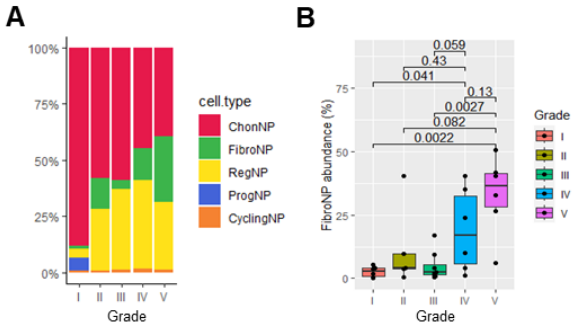

**Supplementary Figure S3. Histological evaluation of nucleus pulposus fibrosis in human disc degeneration**. Various staining were performed on NP sections representing non-degenerative (nNP) or degenerative (dNP) condition: Hematoxylin and eosin staining (H&E); Multichromatic staining of Fast green/Alcian blue/Safranin O/Tartrazine (FAST); Polarized microscopy of Sirius red (PSR); immunofluorescence staining of fibrotic collagen I (COL1, green) and collagen III (COL3, green), and hyaline matrix components of aggrecan (ACAN, green) and collagen II (COL2, red). Arrow head: round-shaped cells; Arrow: spindle-shaped cells.

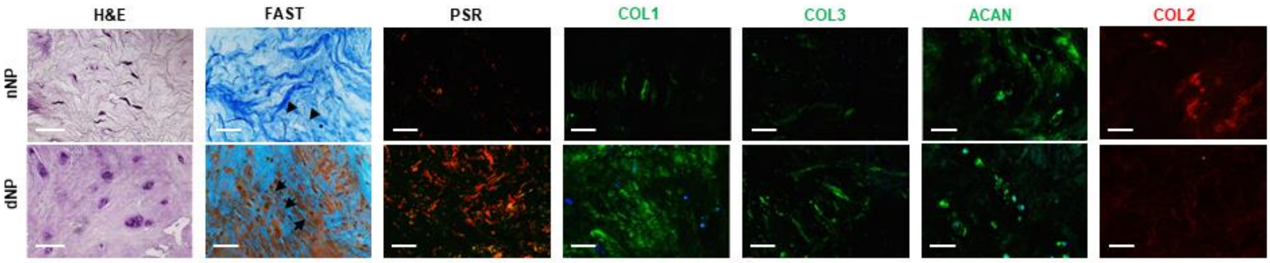


**Supplementary Figure S4. Flow cytometry analysis of COL1$^{+}$ NP cells.** (A) Representative COL1 flow cytometry of freshly isolated NP cells from non-degenerative scoliotic (nNP) and degenerative and herniated (dNP) discs. (B) Quantification of COL1$^{+}$ cells (n=3). Two-tailed unpaired *t*-tests.

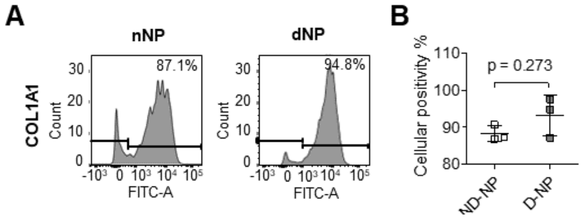

**Supplementary Figure S5. Characterization of myofibroblastic phenotype in NP cells culture. (A)** Representative immunofluorescence of cultivated primary cells from scoliotic (nNP) and degenerative (dNP) human NP tissues. Scale bar: 50µm. **(B)** RT-qPCR of gene markers of myofibroblast and NPCs. *p < 0.05, **p < 0.01 by one sample *t*-tests.

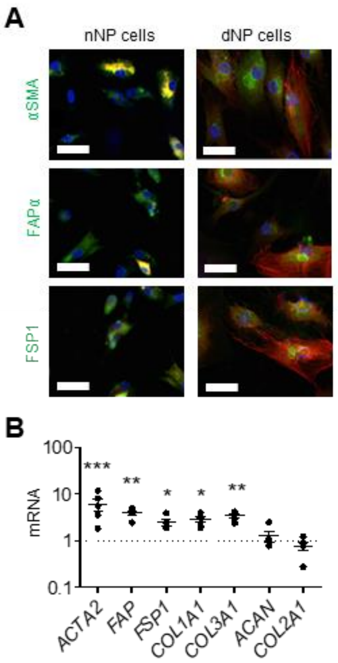

**Supplementary Figure S6. Analysis of putative fibrocyte marker CD34.** UMAP distribution and immunodetection of co-expression for *CD34* with *COL1A1* (**A**) and *ACTA2* (**B**) in disc cells. Immunostaining was conducted on human NP tissue sections from non-degenerative scoliotic (nNP) or degenerated and herniated IVDs (dNP). CD34 was in green, and COL1/αSMA in red. Scale bar: 50µm. Insert: representative cells micrograph showing positive signals; scale bar: 12µm.

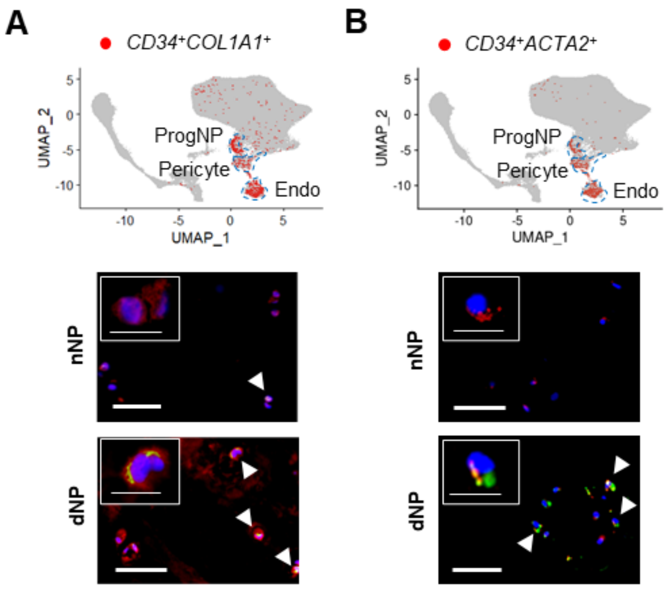

**Supplementary Figure S7. GO (gene ontology) functions of immunocyte- and NPC-clustered fibrocytes.** (A) from 47 positively expressed DEGs and (B) from 236 negatively expressed DEGs of immunocyte-clustered fibrocytes; (C) from 43 positively expressed DEGs of NPC-clustered fibrocytes. Fib-M/N/T/G: fibrocyte distributed in macrophage, neutrophil, T cell and G-MDSC clusters; fib-cNP/fNP/rNP: fibrocyte distributed in ChonNP, FibroNP and RegNP clusters.

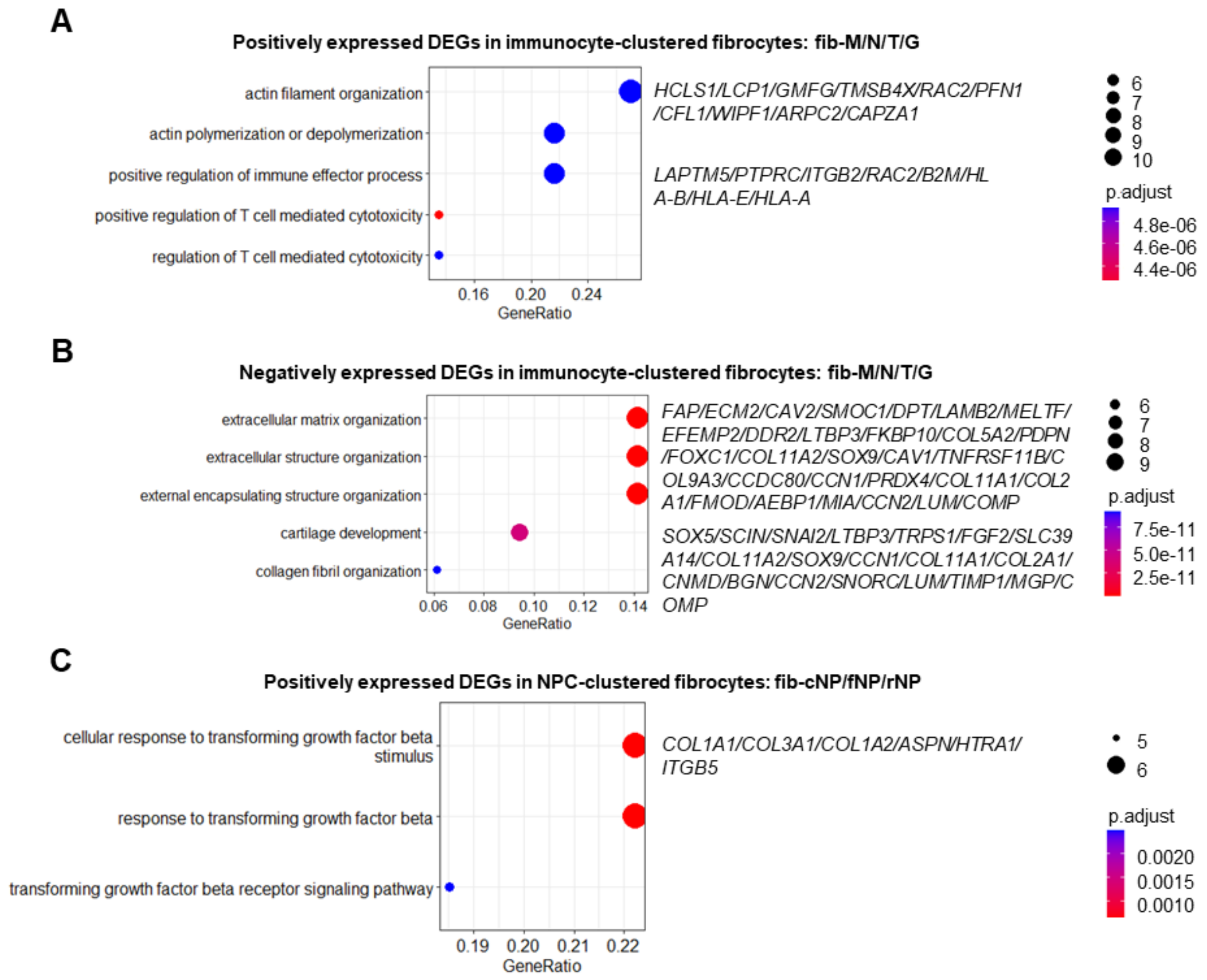

**Supplementary Figure S8. Bubble plot from CellChat analysis showing the molecular pattern of interaction between disc fibrocytes and the NPC clusters.**

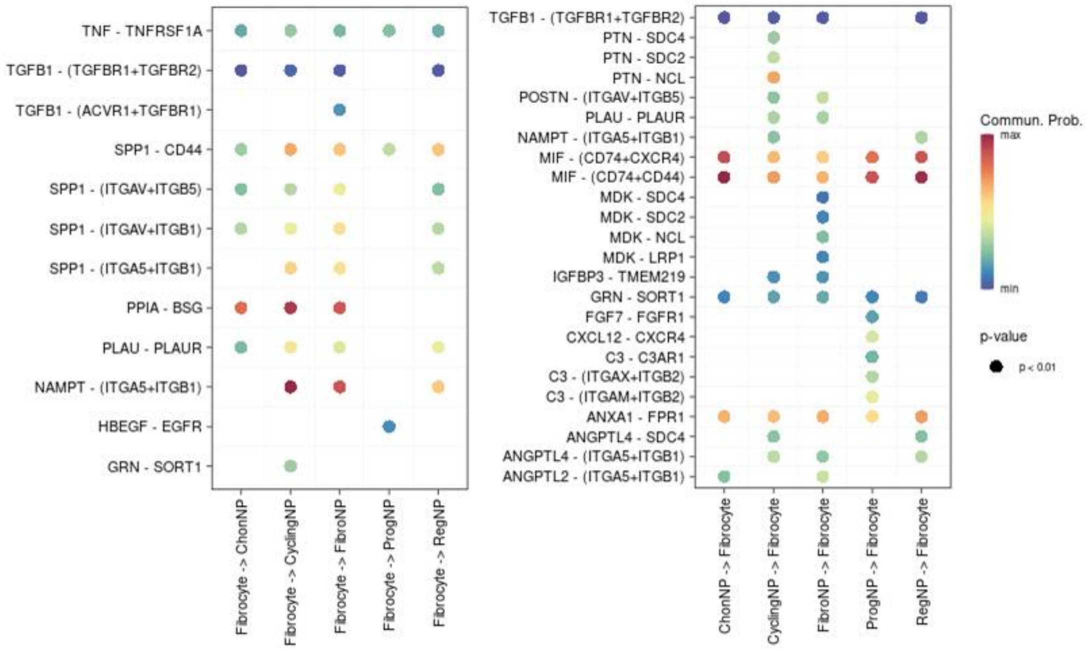

**Supplementary Figure S9. Immunodetection of monocytic cells by GFP. (A)** Bone marrow stain; (B) Healthy discs of CD11b-DTR mice. Non-punctured discs serve as sham control and negative to GFP staining; (C) Monocytic fibrocyte presence in annulus fibrosus (AF). Scale bar: 50µm. dpp: days post-puncture; wpp: weeks post-puncture.

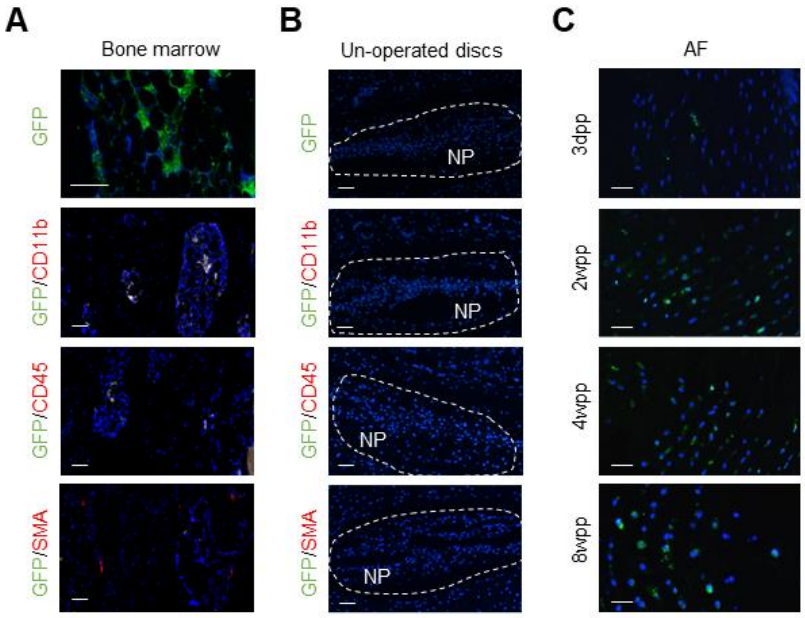

**Supplementary Figure S10. Monocyte depletion in CD11b-DTR mice. (A)** Representative flow cytometry of CD11b$^{+}$ monocyte in peripheral blood. Isotype control was shown in blue. (B) Representative micrographs for GFP stain in punctured discs. DT injection was conducted on CD11b-DTR mice 3 days prior to disc puncture and 4 weeks later at dosage of 10ng/g. Scale bar: 50μm. wpp: weeks post-puncture.

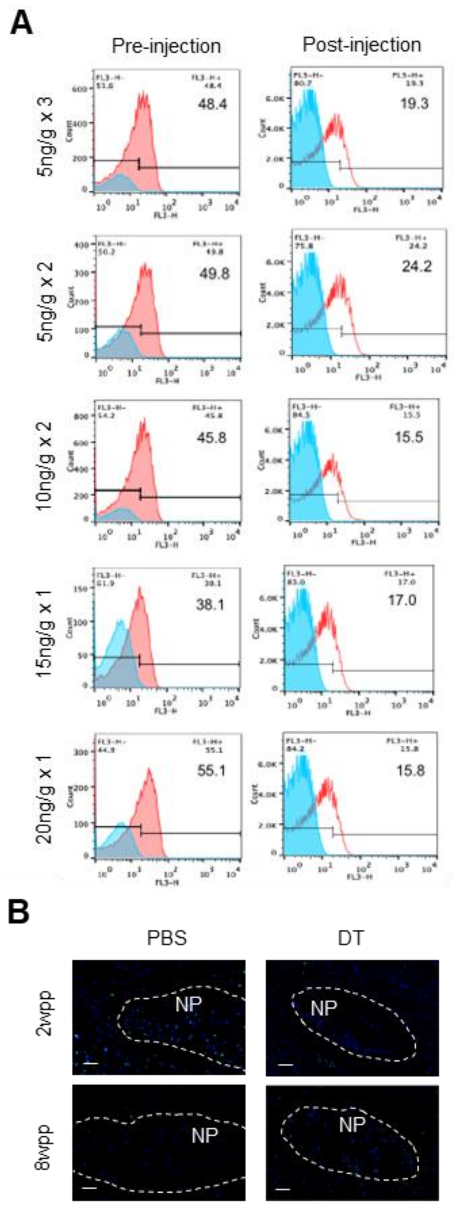

**Supplementary Table S1. Summarization of disc cells atlas and conventional cellular markers.**

Four published single cell RNA sequencing analysis using human disc/NP tissues were included.

| Ref | Annotation | Defined markers | Additional information |
|---|---|---|---|
| | C1 chonNPC | *MMP3, PDE4B,* ***CXCL8****, TNFRSF11B, MGP, ADPRHL1, FGF2, BMP2, C11orf96, SOD2,* ***CXCL2****, GPX3, CLU* | Inflammatory process |
| | C2 chonNPC | *PRG4, SPARC, MSMO1, CYTL1,* ***COL2A1****, ABI3BP, S100A2,* ***VCAN****, TIMP1, HAPLN1* | Cholesterol biosynthetic process and maintenance of the structure |
| | C3 chonNPC | ***COL12A1****, SAA1,* ***MMP13****, TNC, CXCL1, CHI3L2,* ***MMP2****, AEBP1, GREM1, IFITM1* | Inflammatory responses and matrix disassembly |
| | C4 chonNPC | *POSTN,* ***COL1A1****,* ***COL3A1****, COL1A2, SPP1,* ***TMSB4X****, LGALS1, MMP13, COL6A3, MMP14* | Fibrous characteristics |
| 7 | Cartilage Progenitor | *STMN1, HIST1H4C, LGALS1, NUSAP1,* ***TOP2A****, HIST1H1B, TUBB, CKAP2, PTTG1, POSTN,* ***CENPF****,* ***BIRC5*** | NP-derived progenitor |
| | Fibrochondrocyte Progenitor | *MYLK,* ***TRIB3****, TCEA1, ATF3, ATF5, SYNE1, GOT1, GDF15, CYR61, DEPP1,* ***CANX*** | Protein folding quality control and ER stress |
| | Homeostatic chondrocytes | ***FOS****,* ***JUN****, ZFP36, RGS16, IER2, ID3, JUNB, ATF3, EGR1, IRF1* | Responsive to stress |
| | Endothelial cells | *HES1, CALCRL, COL4A1, SPARCL1,* ***PECAM1****, IFI27, PLVAP, AQP1, COL15A1, LAMA4,* ***CD34*** | |
| | Macrophage | *CCL3L1, CCL4L2, CCL4, IL1B,* ***CD74****, CXCL3, CXCL8, HLA-DRA, CCL3, TNF,* ***LAPTM****, TYROBP* | |
| 14 | NP progenitor | *PLA2G2A, FBLN1, SERPINF1, CFD, PTGDS, WISP2, IGFBP6, GSN, MMP2, C1R,* ***PDGFRA****,* ***PAX1****,* ***ANGPT1****, PRG4,* ***PRRX1*** | Bone morphogenesis, connective tissue development |

| | | | |
|---|---|---|---|
| | Stromal cells | *ACTA2, MYL9, RGS5, MYH11, SYNPO2, STEAP4, NOTCH3, MCAM, ADIRF, ADIRF*, ***FOXC2***, ***GJA1***, ***HES4*** | |
| | Stromal cells-Fibroblast | *CEMIP, AKR1C1, MGP, COMP, DNER, MELTF* | |
| | Stromal cells-Neurogenic | *SOX2, NGFR, NCMAP, CLDN19* | |
| | Stromal cells-Osteogenic | *RUNX2, DLX5, SP7, BGLAP, MMP11* | |
| | Pericyte | *ITGB1, CARMN, ID3, C1QTNF1, MYL6, MAP3K20, TNS1, LGI4, ABCC9*, ***ACTA2***, ***TAGLN***, ***MCAM*** | |
| | Chon1 (C1/2) | ***CYTL1***, ***IBSP***, *CRYAB, APOD, IL17B, RBP4, C2orf40, CHAD, SPP1, HSPA6* | Regulatory chondrocytes secreting growth factors, AF and CEP chondrocytes |
| | Chon2 (C3/4) | *NEAT1, MALAT1, DST, COL11A1*, ***COL2A1***, *FN1, ACAN, FMOD*, ***COMP***, *SLC5A3*, ***CCNL1***, ***WSB1*** | Homeostatic chondrocytes, ECM homeostasis and circadian rhythm |
| | Chon3 (C5/6) | *CHRDL2, CAPS, NDUFA4L2, ABI3BP*, ***CNMD***, ***CHI3L1***, *MT1E, MT1X, VCAN, CRISPLD1*, ***PRG4***, ***KLF2***, ***COL5A1***, ***EPYC*** | Effector chondrocytes, metabolic active |
| | Notochord NPC | ***KRT19***, ***CA3***, *EPYC, ENPP2, CD24, CTHRC1, ACTC1, PCSK2, RAPGEF5, RAB3B*, ***KRT8***, ***TBXT*** | |
| | Endothelial cells | *CD74, IFI27*, ***PECAM1***, *RAMP2, GNG11, CALCRL*, ***VWF***, *EGFL7, STC1, HLA-E*, ***CD34***, ***CDH5***, ***ERG*** | |
| | Blood cells | *LYZ, S100A9, S100A8, CXCL8, SRGN, AC020656.1, MPO, CCL3, TYROBP, AZU1* | |

| | | | |
|---|---|---|---|
| 12 | Effector NPC | *MSMO1, HMGCS1, INSIG1* | Metabolic process, Positive regulation of ECM assembly |
| | Hypertrophic chonNPC | *FRZB, DKK1, EGR1* | Programmed cell death; ECM disassembly |
| | Adhesion NPC | *FN1, CRTAC1, FMOD* | Cell migration, cell–matrix adhesion. |
| | FibroNPC | *COL1A1, COL3A1, MMP2, COL6A1, TGFB* | Fibrosis related |
| | Homeostatic NPC | *RPS29, RPS21, RPL31* | Cellular homeostasis |
| | Regulatory NPC | *CHI3L1, NFKB, CXCL2, CXCL3, IL6* | Cellular responses to inflammation and endogenous stimuli |
| | NK cells | *CD94* | |
| | Macrophage | *CD163* | |
| | T cell | *TRAC* | |
| | G-GMP | *MS4A3, MPO, ELANE* | |
| | Neutrophil | *FCGR3B, HLADR* | |
| | G-MDSC | *ITGAM, OLR1, ARG1, CD45* | Immunopression, ROS production |
| 11 | Endothelial progenitor | *PECAM1, CDH5, CD34, KDR* | |
| | Erythrocytes | *HBA1, HBB* | |
| | Effector NP cells, EffectorNP | *COL2A1, SPARC, CLEC3A, COL3A1, FMOD, CYTL1* | ECM organization, cartilage development |
| | Homeostasis NP cells, HomNP | *MT1G, EMP1, HSPB1, S100A2, RGCC, ANXA1* | Responsive to stress, detoxification of inorganic compound |
| | Hypertrophic NP cells, HTNP | *MMP3, CXCL2, SOD2, C11orf96, BMP2, SERPINE2* | Cellular response to external stimulus, inflammatory reaction |
| | NP progenitor cells, NPPC | *TMSB4X, NEAT1, XIST, MALAT1* | Chondrocyte differentiation, |

| | | | |
|---|---|---|---|
| | | | extracellular structure organization, ossification |
| | CD24+ progenitor | *CD24, KRT19, KRT8, GJA1, LGALS3, APOE, CA3* | Protein synthesis, cellular pluripotency |
| | MK167+ progenitor | *MK167, CENPF, STMN1, POSTN, TUBB, TUBA1B, COL1A1* | Epithelial-mesenchymal transition (EMT), inflammatory response |
| | AF cells | *COL1A1, CRTAC1, ASPN, MMP2* | |

**Supplementary Table S2. Fibrocyte count in NP cells clusters.** Either *ITGAM*$^{+}$*COL1A1*$^{+}$ or *PTPRC*$^{+}$*COL1A1*$^{+}$ cells are regarded as disc fibrocytes, and their number in NP cells clusters (as described in Fig. 1) were count.

| **NP cells clusters** | ***ITGAM*$^{+}$*COL1A1*$^{+}$** | | ***PTPRC*$^{+}$*COL1A1*$^{+}$** | | ***Disc fibrocytes*** | |
|---|---|---|---|---|---|---|
| | **Count** | **%** | **Count** | **%** | **Count** | **%** |
| ChonNP | 38 | 3.47 | 54 | 3.11 | 89 | 4.16 |
| FibroNP | 97 | 8.87 | 69 | 3.97 | 160 | 7.48 |
| RegNP | 36 | 3.29 | 30 | 1.73 | 64 | 2.99 |
| ProgNP | 0 | 0.00 | 3 | 0.17 | 3 | 0.14 |
| CyclingNP | 4 | 0.37 | 6 | 0.35 | 9 | 0.42 |
| Neutrophil | 411 | 37.57 | 647 | 37.23 | 732 | 34.24 |
| G-MDSC | 21 | 1.92 | 52 | 2.99 | 63 | 2.95 |
| Macrophage | 462 | 42.23 | 629 | 36.19 | 756 | 35.36 |
| T cell | 16 | 1.46 | 238 | 13.69 | 243 | 11.37 |
| Endothelial cell | 4 | 0.37 | 3 | 0.17 | 7 | 0.33 |
| Pericyte | 4 | 0.37 | 6 | 0.35 | 10 | 0.47 |
| Erythrocyte | 1 | 0.09 | 1 | 0.06 | 2 | 0.09 |

**Supplementary Table S3. Safety evaluation of DT injection in CD11b-DTR mice.** Depletion efficiency was calculated based on number of CD11b+ cells in peripheral blood. Three animals were included for each condition. A: >1 living mice at week 8; D: all three mice dead at week 8.

| **Dosage** | **Frequency** | **Time frame/ weeks** | | | | | **Depletion efficiency** |
|---|---|---|---|---|---|---|---|
| | | **1st** | **3rd** | **5th** | **6th** | **8th** | |
| 5ng/g | 3 | √ | √ | √ | | A | 55.80% |
| | 2 | √ | | √ | | A | 45.78% |
| 10ng/g | 3 | √ | √ | √ | | D | N/A |
| | 2 | √ | | √ | | A | 65% |
| 15ng/g | 2 | √ | | √ | | D | N/A |
| | 1 | √ | | | | A | 55.46% |
| 20ng/g | 2 | √ | | √ | | D | N/A |
| | 1 | √ | | | | A | 68.50% |
| 25ng/g | 2 | √ | | √ | | D | N/A |
| | 1 | √ | | | | D | N/A |

**Supplementary Table S4. Demographics of IVD donors**. Grade of disc degeneration was determined by MRI according to Pfirrmann Scale. F, female; M, male; AIS, adolescent idiopathic scoliosis; DDD, disc degeneration disease; ND, not determined.

| Sample ID | Gender | Age (Years) | Level | Pathology | Disc grade |
|---|---|---|---|---|---|
| 1 | F | 13 | L2-3 | AIS | ND |
| 2 | F | 13 | L2-3 | AIS | ND |
| 3 | M | 15 | L2-3 | AIS | ND |
| 4 | F | 15 | L2-3 | AIS | ND |
| 5 | F | 16 | L1-2 | AIS | ND |
| 6 | F | 15 | L2-3 | AIS | ND |
| 7 | M | 28 | L5-S1 | DDD | IV |
| 8 | M | 40 | L3-L4 | DDD | III |
| 9 | F | 42 | L4-L5 | DDD | IV |
| 10 | F | 47 | L3-L4 | DDD | V |
| 11 | F | 56 | L4-L5 | DDD | III |
| 12 | M | 58 | L3-L4 | DDD | V |
| 13 | M | 59 | L4-L5 | DDD | IV |
| 14 | F | 45 | L4-L5 | DDD | IV |
| 15 | M | 71 | L4-L5 | DDD | IV |

**Supplementary Table S5. Antibodies used for immunofluorescence and flow cytometry (FC) analysis.**

| Product Cat# | Antigen | Host | Conjugation | Application |
|---|---|---|---|---|
| EPR14664 | Aggrecan | Rabbit | N/A | ACAN in human NP (Supplementary Fig. S1) |
| Ab34710 | Collagen I | Rabbit | N/A | COL1 in human NP (Supplementary Fig. S1); FC for COL1 (Supplementary Fig. S4); Co-stain with CD34 in Cd11b-DTR mice IVDs |
| Ab88147 | Collagen I | Mouse | N/A | FC for COL1 (Fig. 2); Co-stain with CD45, αSMA in human NP (Fig. 2) |
| EPR12268 | Collagen II | Rabbit | N/A | COL2 in human NP (Supplementary Fig. A1) |
| Ab7778 | Collagen III | Rabbit | N/A | COL3 in human NP (Supplementary Fig. S1) |
| EPR5368 | αSMA | Rabbit | N/A | FC for αSMA (Fig. 1); αSMA in human NP (Fig. 1); co-stain with GFP in CD11b-DTR mice IVDs (Fig. 3) |
| Ab7817 | αSMA | Mouse | N/A | Co-stain with CD45 in human NP (Fig. 2) |
| EPR20021 | Fap-α | Rabbit | N/A | FC for FAPα (Fig. 1); FAPα in human NP (Fig. 1); |
| EPR2761-2 | Fsp1 | Rabbit | N/A | FC for FSP1 (Fig. 1); FSP1 in human NP (Fig. 1); |
| Ab23910 | CD45 | Rat | N/A | Co-stain with COL1 and αSMA in CD11b-DTR mice IVDs (Fig. 4) |
| Ab10558 | CD45 | Rabbit | N/A | Co-stain with COL1 in human NP (Fig. 2), and GFP in CD11b-DTR mice IVDs (Fig. 3) |
| 9B10D4 | CD34 | Mouse | N/A | Co-stain with COL1 and αSMA in human NP (Supplementary Fig. S6) |
| Ab184308 | CD11b | Rabbit | N/A | Co-stain of CD11b with GFP in CD11b-DTR mice IVDs (Fig. 3); FC for CD11b in CD11b-DTR mice IVDs (Supplementary Fig. S9) |
| Ab5450 | GFP | Goat | N/A | GFP in CD11b-DTR mice IVDs (Fig. 3) |

| | | | | |
|---|---|---|---|---|
| Ab150077 | Rabbit IgG H&L | Goat | Alexa Fluor® 488 | Immunostaining of COL1/3, ACAN, αSMA, FAPα (Supplementary Fig. S1 and Fig. 1); FC for αSMA, FAPα, FSP1, COL1 and CD45 in human NP cells (Fig. 1 & 2, Supplementary Fig. S4); co-stain of CD45 with COL1 and αSMA in human NP (Fig. 2); co-stain of COL1 and αSMA with CD45 in CD11b-DTR mice (Fig. 4) |
| Ab150080 | Rabbit IgG H&L | Goat | Alexa Fluor® 594 | Immunostaining of COL2, FSP1 in human NP (Supplementary Fig. S1 & Fig. 1) and COL1 in CD11b-DTR mice IVDs (Fig. 4); FC for αSMA, Fap-α and Fsp1 (Fig. 1); Co-stain of CD45 with COL1 (Fig. 2), COL1 and αSMA with CD34 in human NP (Supplementary Figu. S6) |
| Ab150116 | Mouse IgG H&L | Goat | Alexa Fluor® 594 | FC for COL1 (Fig. 2); Co-stain of COL1 and αSMA with CD45 in human NP (Fig. 2) |
| Ab150160 | Rat IgG H&L | Goat | Alexa Fluor® 594 | Co-stain of CD45 with COL1 and αSMA in CD11b-DTR mice IVDs (Fig. 4) |
| Ab150113 | Mouse IgG H&L | Goat | Alexa Fluor® 488 | Co-stain of COL1 and αSMA with CD34 in human NP (Supplementary Fig. S6) |
| Ab150129 | Goat IgG H&L | Donkey | Alexa Fluor® 488 | GFP and co-stain with CD11b, CD45, αSMA in CD11b-DTR mice IVDs (Fig. 3) |
| Ab175470 | Rabbit IgG H&L | Donkey | Alexa Fluor® 568 | Co-stain of CD11b, CD45, αSMA with GFP in CD11b-DTR mice IVDs (Fig. 3) |

**Supplementary Table S6.** Primers for real-time PCR.

| Gene name | Forward | Reverse |
|---|---|---|
| *ACTA2* | CTATGAGGGCTATGCCTTGCC | GCTCAGCAGTAGTAACGAAGGA |
| *FAP* | ATGAGCTTCCTCGTCCAATTCA | AGACCACCAGAGAGCATATTTTG |
| *S100A4* | GATGAGCAACTTGGACAGCAA | CTGGGCTGCTTATCTGGGAAG |
| *COL1A1* | ATCAACCGGAGGAATTTCCGT | CACCAGGACGACCAGGTTTTC |
| *COL3A1* | GCCAAATATGTGTCTGTGACTCA | GGGCGAGTAGGAGCAGTTG |
| *COL2A1* | TGGACGATCAGGCGAAACC | GCTGCGGATGCTCTCAATCT |
| *ACAN* | ACTCTGGGTTTTCGTGACTCT | ACACTCAGCGAGTTGTCATGG |
| *GAPDH* | GGAGCGAGATCCCTCCAAAAT | GGCTGTTGTCATACTTCTCATGG |

**Supplementary methods**

*Disc height index calculation*

Based on the X-rays for coccygeal IVDs of Cd11b-DTR mice, the lengths of the discs and adjacent vertebral bodies were measured with Image J (Version 1.42, National Institutes of Health). Disc height index (DHI) was calculated according to the formula: DHI = the disc height (averaging three measurements of the heights of lateral and middle portions of the disc)/ the vertebral body height (averaging three measurements of the heights of the lateral and middle adjacent caudal vertebral body), and. Level C6/7 of tail disc was un-operated and further used as reference for DHI calculation.
